# Mitochondrial Genome Variants and Nuclear Mitochondrial DNA Segments in 7331 Individuals from NyuWa and 1KGP

**DOI:** 10.1101/2025.06.21.660892

**Authors:** Yuanxin Wang, Jiajia Wang, Yanyan Li, Peng Zhang, Zhonglong Wang, Shuai Liu, Yiwei Niu, Yirong Shi, Sijia Zhang, Tingrui Song, Tao Xu, Shunmin He

## Abstract

Dysfunctional mitochondria are implicated in various diseases, while little is known about comprehensive characterization of mitochondrial DNA (mtDNA) in Chinese population. Here, we conducted a systematic analysis of mtDNA from 7331 samples, comprising 4129 Chinese samples (NyuWa) and 3202 samples from the 1000 Genomes Project (1KGP). We identified 7216 distinct high-quality mtDNA variants and classified them into 22 macrohaplogroups, and detected 1466 distinct nuclear mitochondrial DNA segments (NUMTs), with 88 mtDNA variants and 642 NUMTs being specific to NyuWa. The genome-wide association analyses revealed that 12 mtDNA variants were significantly correlated with 199 nuclear DNA variants. Our findings revealed that all individuals in both NyuWa and 1KGP harbored common NUMTs, and one-fifth possessed ultra-rare NUMTs, which tended to insert into nuclear gene regions. Compared to 1KGP, significant enrichment of nuclear breakpoints in long interspersed nuclear elements (LINEs) was observed for rare NUMTs in NyuWa. Overall, this study represents the first comprehensive profiling the landscape of Chinese NUMTs and offers the most extensive resource of Chinese mtDNA variants and NUMTs based on high-depth WGS to date, providing valuable reference resources for genetic research on mtDNA-related diseases.

## Introduction

Mitochondria are essential organelles that provide the majority of energy for the organisms through oxidative phosphorylation in mammalian cells, and have genetic material that has a strong selective effect on the nuclear genome [1,2]. The mitochondrial DNA (mtDNA) of human is a circular double-stranded DNA molecule containing 16,569 base pair, consisting of 22 transfer RNA (tRNA) genes, 2 ribosomal RNA (rRNA) genes, 13 protein-coding genes, and other non-genic regions (D-loop and intergenic regions) [3]. mtDNAs are polyploid, since each cell contains thousands of mitochondria and each mitochondria contains multiple copies of mtDNA [4,5].

mtDNA has a high mutation rate due to the lack of DNA damage repair, high oxidation environment, increased replication times, maternal inheritance and lack of recombination [4,6–8]. The higher mtDNA mutation rate is associated with regional mtDNA variation reflective of prehistoric human migration patterns [9]. Studies have shown that variation in the intracellular percentage of normal and mutant mtDNA (heteroplasmy) can be associated with phenotypic heterogeneity in mtDNA diseases[10–12]. The heteroplasmic level can accumulate gradually with age, and when it exceeds a critical threshold, a corresponding phenotype may occur [13,14]. mtDNA variants are reported to be associated with a variety of complex diseases, like neurological diseases [15–17], metabolic related diseases [6,18–21], and cancers [22]. NUMTs refer to cytoplasmic mtDNA fragments that are transferred into the nuclear genome. More NUMT sequences of different sizes and lengths in the diverse number of eukaryotes have been detected as whole genome sequencing (WGS) of different organisms accumulates. NUMT is an ongoing phenomenon that can be transmitted from parents to offspring and occur approximately once every 10^4^ human births [23]. NUMTs are observed to have non-random distribution and it may disrupt protein-coding genes depending on the position of the insertion [23,24]. The transposition of NUMTs into the genome has been associated with human diseases, such as mucolipidosis IV [25], Pallister-Hall syndrome [26], Usher syndrome type IC [27], plasma factor VII deficiency [28] and cancers [23,29].

Population-specific genomics are fundamental for genetic disease research and precision medicine. Mitochondria play important roles in biological processes, including apoptosis [30–32], lifespan modulation [32], cytoplasmic calcium buffering [33], innate immunity [34] and cell cycle [35,36]. The development of WGS technology has enabled us to analyze genetic variation, NUMTs and their functions in large-scale populations from different lineages. The representative population-scale mtDNA studies, such 1KGP and gnomAD, have primarily been based on individuals of European descent, and there are few studies on mtDNA variants and NUMTs based on deep WGS data involving East Asian populations, especially Chinese populations [23,37–40]. We know that as humans migrated, new variants defined geographically located ‘macro-haplogroups’, which differ between continents [41]. Besides, research has shown that major differences in the frequency and distribution of NUMTs between different ethnic groups, with the most obvious difference in East Asia[23]. Our previous works [42–44] have demonstrated the superiority of the Chinese-specific cohort, NyuWa, in discovering novel functional rare variants. Considering a certain degree of population heterogeneity, it is necessary to construct a genetic map of mtDNA variants and NUMTs in Chinese people to fill the missing diversity worldwide. In view of this, we conducted a systematic analysis of mitochondrial DNA comprehensive characterization from 7331 samples focusing on the Chinese population, which were also utilized to profile the landscape of the Chinese NUMTs. We provided the largest mtDNA variant and NUMTs resource of Chinese population based on high-depth WGS data so far.

## Results

### 7216 high-quality mtDNA variants and 824 distinct NUMTs

We conducted a systematic analysis of mtDNA comprehensive characterization from two cohorts: NyuWa dataset consisting of 4129 Chinese individuals and the 1KGP dataset consisting of 3202 samples. The NyuWa samples were collected from 24 administrative divisions in China, including 17 provinces, 3 autonomous regions, and 4 municipalities directly under the central government. The majority of these samples were from East, North, and South China, with significant representation from Shanghai, Guangdong, and Beijing. The ethnic composition is predominantly Han Chinese, as national minorities are geographically clustered and not well-represented in the sampling areas. The sex of the samples was determined based on the genomic coverage of sex chromosomes, and there were 1787 females and 2342 males.

As the WGS data contain information of mtDNA sequence, we identified mtDNA variants using the mitochondrial variant calling pipeline [37] (Table S1, Methods) and the non-reference NUMTs using NUMTs-detection method [23] in 7331 WGS samples from NyuWa and 1KGP. In NyuWa and 1KGP cohort, the median depth of nDNA was 30X and 32X (Figure S1A), respectively. The median depth of mtDNA was ∼3000X and ∼12,000X (Figure S1B), respectively. To filter out potential duplicate and contaminated samples, three methods were used including mtDNA copy number, assessment of nDNA contamination by VerifyBamID2 and assessment of mtDNA contamination by Haplocheck [45] (Figure S1C). As a result, 7266 samples remained for mtDNA variants analysis, and a weak correlation between nuclear and mtDNA depth was observed (Figure S1D). On the basis of removing contaminated samples and duplicate data, we removed samples according to insert size, 7324 samples were used to profile the landscape of the NUMTs.

Using at least 5 discordant read pairs filtering to ensure the stringency, 120,400 NUMTs that are not present in the reference sequence were detected, and each sample identified a mean of 16.44 NUMTs (s.d. = 11.91) (Figure 1A). Grouping NUMT clusters from multiple samples, we observed 1466 distinct NUMTs, which ranged from 5bp to 16,568bp in length, with a median of 119bp, or a mean of 881.7bp (s.d. = 2871.41) (Figure 1B). Compared with the NUMTs of 100,000 Genomes Project (63.2% of NUMTs were less than 200 bp and 77.8% were less than 500 bp in size), there was a greater proportion of short insertions (88.51% of NUMTs were less than 400 bp and 89.15% were less than 500 bp in size)(Figure 1B). All NUMTs was divided into four categories based on population frequency: common (frequency (F) ≥ 1%), rare (0.1% ≤ F < 1%), ultra-rare (F < 0.1%) and private (detected in only one person). The ‘common’ category comprises 6.21% of the total and the ‘rare’ category constitutes 9.00%. 1243 NUMTs were ultra-rare (contained class private), which encompasses the majority of the dataset at 84.79% (Figure 1C).

**Figure 1.**
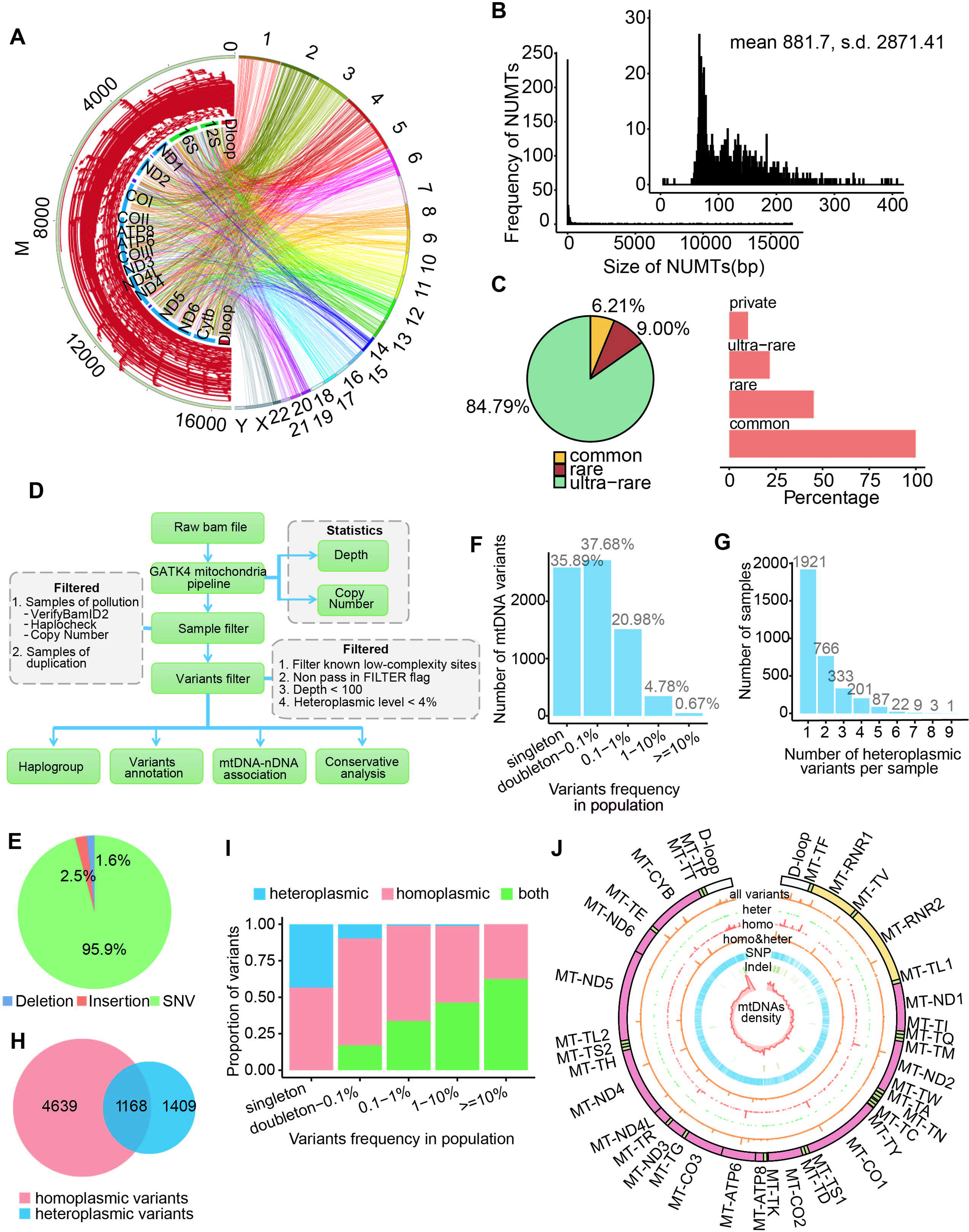
Characteristics of mtDNA variants and NUMTs in this study. **A**. Circos plot of all NUMTs. Left, size and location of NUMTs on mtDNA. Right, chromosomal locations of NUMT insertion. **B**. Size distribution of NUMTs. NUMTs less than 400 bp in length are shown in the inset. **C**. Left, the proportion of NUMTs by population frequency. Right, proportion of samples carrying NUMTs by population frequency. **D.** Pipeline of mtDNA variants calling and filtering. **E.** Variant type (SNVs, deletions, insertions) composition of distinct high-quality mtDNA variants in this study. **F.** Number of high-quality mtDNA variants detected at different variant frequencies in this study. **G.** Number of heteroplasmic variants per sample in this study. **H.** Number of homoplasmic, heteroplasmic, and their overlap high-quality mtDNA variants in this study. **I.** Proportion of variants (homoplasmic, heteroplasmic, and both) at different population frequencies. **J.** The mtDNA variant spectrum of this study. From outer circle to inner circle: mitochondrial genome, mtDNA variants, heteroplasmic variants, homoplasmic variants, homoplasmic and heteroplasmic variants, SNPs, Indels, and mtDNA variants density.

NUMTs can lead to false-positive variant calls at low variant allele frequencies (VAFs) and may also result in misinterpretation of genuinely homoplasmic variants as heteroplasmic. To reduce false positives caused by NUMTs, we filtered out the samples with low mtDNA copy number which may prone to cause NUMTs misalignment [37], and the depth ratio distribution of nDNA to mtDNA was less than 0.04 (4/100) in all samples (Figure 1D, Figure S1E). That is, if the proportion of reads supporting a variant at the locus exceeded 0.04 (4/104), then the locus was more likely to be a mtDNA variant than NUMTs. Based on this, in order to obtain a high-confidence variant set, we filtered out variants with VAF < 0.1, and the result showed each sample contains at least 2 mtDNA variants, with the maximum number of 101 variants and a median of 36 variants per sample in NyuWa and 1KGP (Figure S1F).

There were 268,746 variants in total at different heteroplasmic levels in NyuWa and 1KGP mtDNA resource, among which 97.8% were homoplasmic variants with VAF >= 0.95, and 2.2% were heteroplasmic variants (VAF: 0.1-0.95) (Figure S2A), and is similar to the proportion of homoplasmic and heteroplasmic variants in the gnomAD dataset (98% were homoplasmic and 2% were heteroplasmic). After deduplication, we finally obtained 7216 distinct high-confidence mtDNA variants from 7,266 individuals. 95.9% of these variants were SNVs (Figure 1E), which were mainly transitions (Figure S2B), especially G>A and T>C (Figure S2C), consistent with the reported mtDNA variation signature in both population-based studies [37,38] and cancer studies [46,47]. As for indels, most were one-base insertions and deletions (Figure S2D). Most of all mtDNA variants were detected at low population frequencies in NyuWa and 1KPG mtDNA resource (Figure 1F), and 47.97% (142/296) of indels were singletons (Figure S2E). 3,343 (46%) samples were detected to contain heteroplasmic variants (Figure 1G), with only one or two heteroplasmic variants in more than 80% of the 3,343 samples. 1,409 (19.53%) mtDNA variants were found only at heteroplasmic levels, 4639 (64.29%) mtDNA variants were observed only at homoplasmy in NyuWa and 1KGP, which is a higher proportion compared to gnomAD (15% heteroplasmic only and 48% homoplasmic only) (Figure 1H), and most insertions were in this category (Figure S2F). In addition, the heteroplasmic variants tended to be at low population frequencies in NyuWa and 1KGP mtDNA resource, while the homoplasmic variants had a relatively wider population frequency spectrum (Figure 1I). 79.77% of variants observed only at heteroplasmic levels were singletons (Figure S2G), suggesting that these variants may be newly generated or harmful to the organism. The map of all mtDNA variants in NyuWa and 1KGP had been shown in Figure 1J, including the distribution of variant population frequency and the density of variants.

In order to sort out the mtDNA variants and NUMTs obtained in this study, we have built NyuWa mtDNA variants resource (NMVR). Users can quickly browse and search to query the mtDNA variant and NUMTs of interest, and obtain relevant information on the variant in NyuWa and other databases like gnomAD, 1KGP, and HelixMTdb.

### Specific mtDNA variants and NUMTs

In order to explore the shared and specific mtDNA variants and NUMTs, we compared NyuWa mtDNA resource with other resources. We found that 4405 (98%, 4405/4493) variants (referring to all mtDNA variants, including homoplasmic variation, heteroplasmic variation, and variation detected at both homoplasmic and heteroplasmic levels) were shared with at least one other database (Figure 2A), indicating the reliability of all mtDNA variants in our call set. The SNVs types of different cohorts were also similar (Figure 2B). Afterwards, we conducted an analysis of variant population frequency correlations using shared variants between NyuWa and other resources. The variant population frequencies in NyuWa were highly correlated with those in other resources (Figure S3A-C). East Asian and South Asian had relatively higher correlation with NyuWa than other subpopulations of 1KGP (Figure 2C-D, S3D-F). SNVs exhibited relatively higher consistency when stratified by variant types (Figure S3G-I). In addition, there were 88 novel variants, named NyuWa-specific mtDNA variants, of which 74% were SNVs (Figure 2E), and more than 50% were in protein-coding genes detected at both homoplasmic and heteroplasmic levels (Figure S3J-K). As expected, the variant frequencies of the NyuWa-specific mtDNA variants were very low. Of the 88 NyuWa-specific variants, 76 variants were singletons (Figure 2F). The position of m.8273CCCCTCT>C, a NyuWa-specific variant observed in 6 individuals, overlaped with m.8273_8281del which was reported to be associated with maternally inherited essential hypertension (MIEH) in China [48].

**Figure 2.**
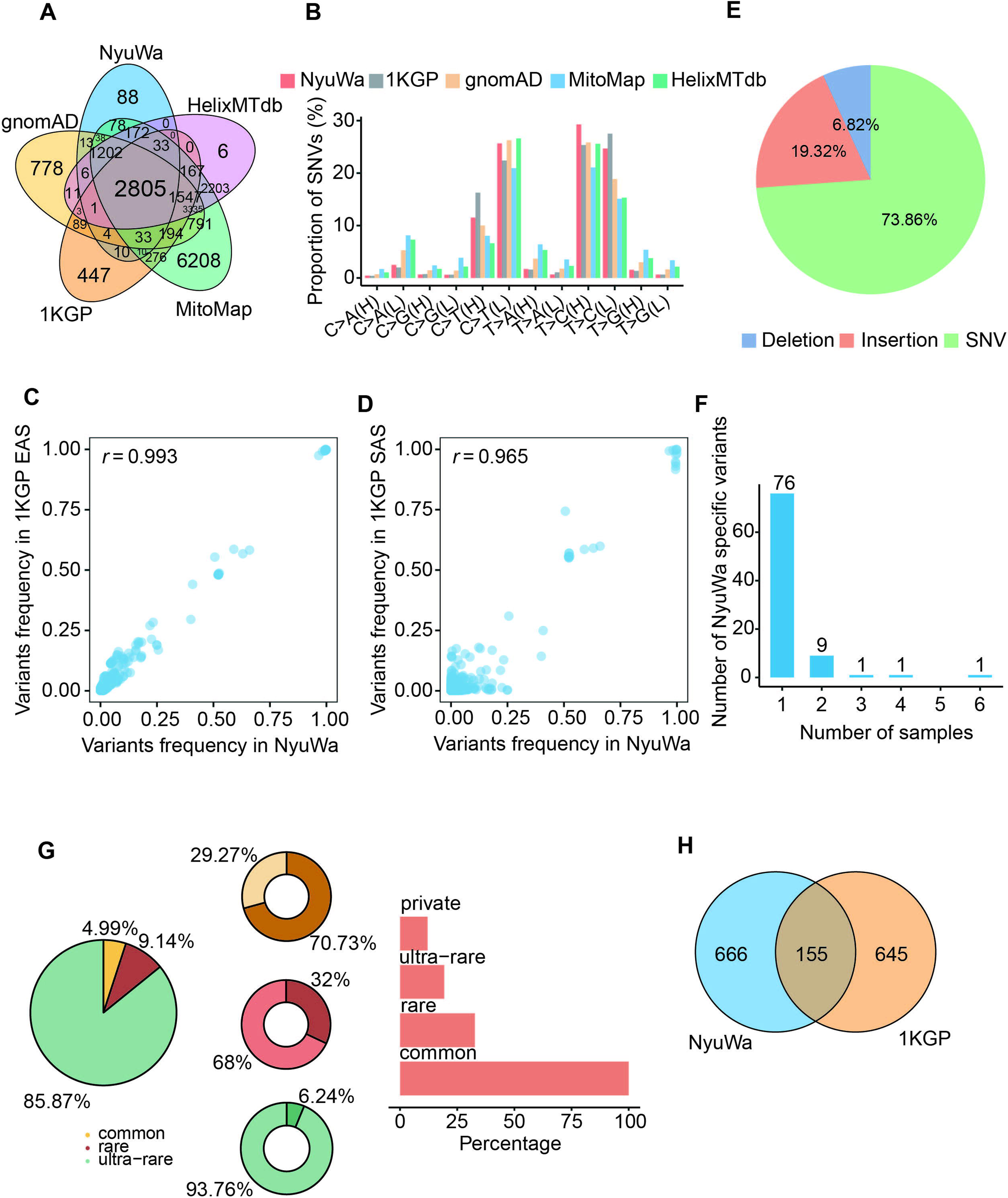
Comparison of mtDNA variants in NyuWa with other resources. **A**. The overlapping variants (referring to all mtDNA variants, including homoplasmic variants, heteroplasmic variants, and variants detected at both homoplasmic and heteroplasmic levels) of NyuWa and other mtDNA variants resources. **B.** The SNVs types of different cohorts. Mitochondrial DNA double strand, H represents the heavy strand and L represents the light strand. **C.** Variant frequencies in NyuWa and 1KGP EAS of this study. EAS, East Asian. Pearson correlation coefficient is shown in the figure. **D.** Variant frequencies in NyuWa and 1KGP SAS of this study. SAS, South Asian. Pearson correlation coefficient is shown in the figure. **E.** Variant type composition of NyuWa-specific mtDNA variants. **F.** Number of samples for NyuWa-specific variants. **G.** Left, the proportion of NUMTs by population frequency. Middle, the proportion of reported (darker colour) and newly (lighter colour) identified NUMTs. Right, proportion of samples carrying NUMTs by population frequency. **H.** Venn diagram of NUMTs distribution in NyuWa and 1KGP.

In NUMTs part, we identified 821 distinct NUMTs in NyuWa. Among these, 705 NUMTs (85.87%) were ultra-rare (contained class private) (Figure 2G). Individuals carrying at least one ultra-rare NUMT accounts for 19.36% of individuals, and 12.11% carried private NUMTs (Figure 2G). Common NUMTs accounts for the smallest proportion, only 5.99%, while the majority of common NUMTs have been reported in other published studies (Figure 2G, Methods) [23,49–52]. On the contrary, only a small number of rare NUMTs were known (Figure 2G). In total, after removing known NUMTs and those overlapping with 1KGP, 642 specific NUMTs were identified among all Chinese NUMTs based on NyuWa Genome resource (Figure 2G-H).

### Functional annotation and pathogenicity of mtDNA variants in NyuWa and 1KGP

To assess the functional impacts of all mtDNA variants in NyuWa and 1KGP, variants were annotated using Variant Effect Predictors (VEP) [53] and Mitochondrial mutation Impact (MitImpact) [54]. 69.71% of high-quality mtDNA variants was found in protein-coding genes (Figure 3A). Most indels (186/296) were in D-loop and intergenic regions, which had weaker effects on biological functions compared with gene regions (Figure S4A). Although the vast majority of the heteroplasmic variants were annotated to D-loop and protein-coding gene regions (Figure S4B), the heteroplasmic variants, especially the variants detected only at heteroplasmic levels, were enriched in rRNA and intergenic regions (Figure 3B). The maximum level of heteroplasmic variants in gene regions was relatively low compared to in D-loop and intergenic regions (Figure 3C), likely because variants with high heteroplasmic level may lead to severe functional impairment. Then we further explored the functional annotation and pathogenicity of mtDNA variants within the Chinese NyuWa dataset alone. Similar trends were observed in both the NyuWa and 1KGP, with the majority of mtDNA variants being located in protein-coding genes (70.38% in NyuWa and 68.96% in 1KGP, Figure S5A and S6A) and lower maximum heteroplasmic levels in gene regions compared to D-loop and intergenic regions (Figure S5C and S6C). Notably, in NyuWa, heteroplasmic variants showed enrichment in D-loop and intergenic regions when compared to 1KGP (Figure S5B and S6B).

**Figure 3.**
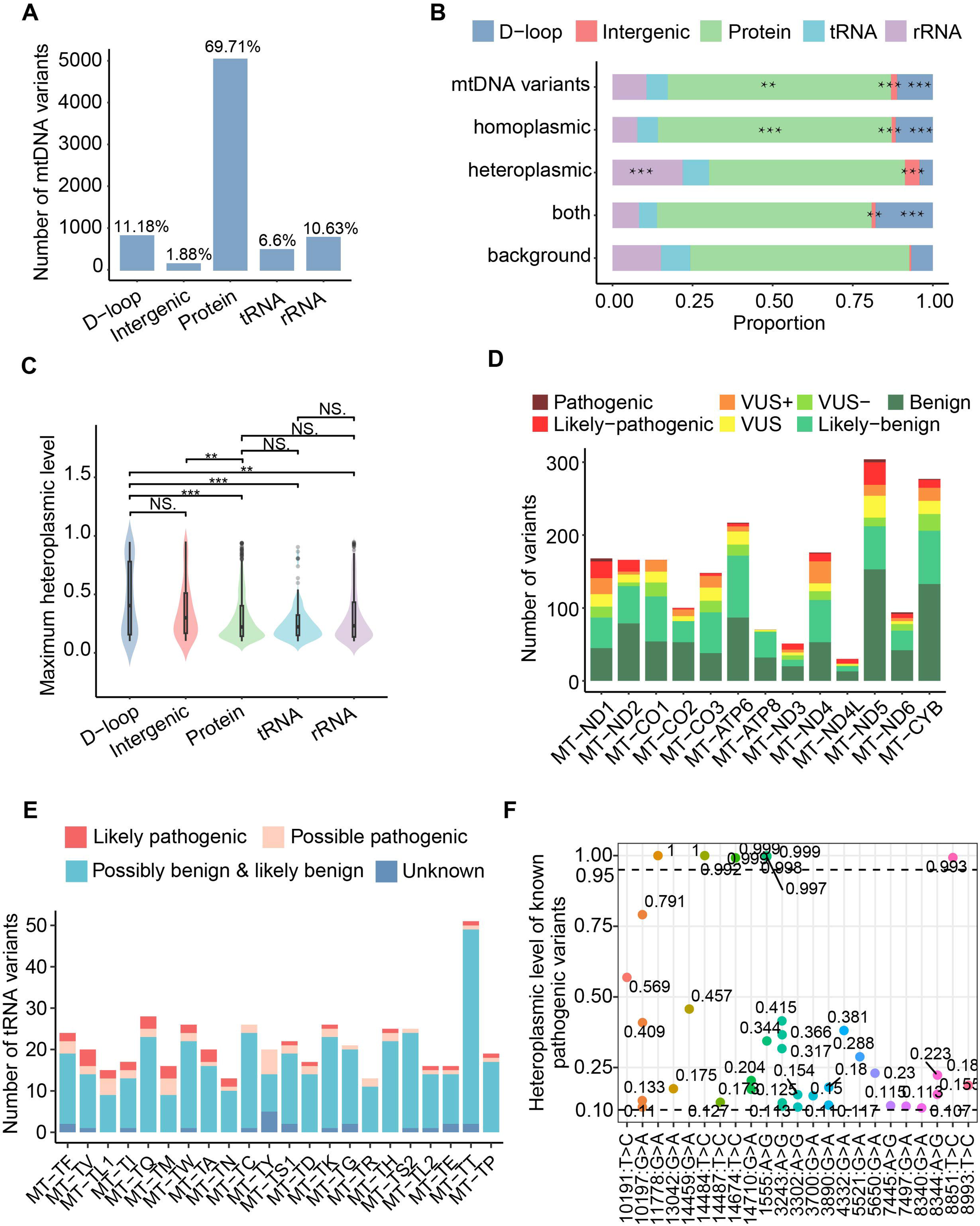
Predicted functions and pathogenicity of mtDNA variants. **A**. Number of mtDNA variants annotated to the mtDNA regions. **B.** Proportion of variant annotations in mtDNA regions. mtDNA variants for total variants. Homoplasmic for variants observed only at homoplasmic levels. Heteroplasmic for variants observed only at heteroplasmic levels. Both for variants observed both at homoplasmic and heteroplasmic levels. Background for the mtDNA genome. ** for *P* <= 0.01, *** for *P* <= 0.001 based on one-tailed Hypergeometric test. **C.** The distribution of the maximum heteroplasmic levels of mtDNA variants in mtDNA regions. *** for *P* <= 0.001, ** for *P* <= 0.01, NS. for no significance based on one-tailed Wilcoxon test. **D.** Number of missense variants in protein-coding genes. APOGEE 2 refers to five pathogenicity classes: benign, likely-benign, VUS, likely-pathogenic, and pathogenic. VUS+ means closer to the likely pathogenic threshold. VUS-means closer to the likely benign threshold. **E.** Number of variants in tRNA genes. The severity of variants was defined by MitoTIP. Likely pathogenic: Variants with high pathogenicity scores, indicating a strong likelihood of causing disease. Possible pathogenic: Variants with moderate pathogenicity scores that may be associated with disease, suggesting a potential but less certain disease-causing effect. Possibly benign & likely benign: Variants with low pathogenicity scores, unlikely to cause disease. Unknown: Variants that do not have a MitoTIP percentile score. **F.** Known pathogenic variants observed in this study along with the heteroplasmic levels. Each dot represents a single individual, and different colors represent different variants.

There were a total of 5030 variants in protein-coding genes, and the homoplasmic levels of variants were widespread in all 13 protein-coding genes (Figure S4C). Of the 5030 variants, 1962 are missense mutations. Pathogenicity annotation was performed using the APOGEE2 meta-predictor from the MitImpact database. Only 13 of the missense mutations were annotated as pathogenic (Figure 3D, Table S2), with 12 identified in the 1KGP and one in the NyuWa (Figure S5D and S6D). The NyuWa pathogenic variant, m.11778G>A in the *MT-ND4* gene, is linked to Leber hereditary optic neuropathy (LHON), a maternally inherited mitochondrial disease causing vision loss [55]. It has been reported that the m.11696G>A mutation may cooperate with other mtDNA variants, raising the penetrance and expressivity of vision loss in Chinese [56].

In addition to protein-coding genes, we also observed the functional disruption of the variants in tRNA genes. The pathogenicity of tRNA associated variants was predicted based on the scoring matrix from MITOMAP [57]. There were 476 variants annotated to tRNA genes, and 31 were assigned as known or likely pathogenic (Figure 3E), of which 26 were identified in 1KGP and 7 were detected in NyuWa (Figure S5E and S6E). Unlike in protein-coding genes, the distribution of heteroplasmic variants in tRNA genes was biased (Figure S4D).

In this cohort, 23 known pathogenic mtDNA variants were present in 39 individuals (Table S3), 9 of whom carried homogenous variation (Figure 3F). m.1555A>G, which was reported to be associated with deaf at homoplasmy, had the highest carrier frequency with 0.0015 (6/4,064) compared to other known pathogenic mtDNA variants in NyuWa cohort (Figure S4E). The prevalence of this pathogenic variant was similar in Chinese population and in multi-population cohort gnomAD (0.0013, 1/763) [37]. Possibly limited by the number of samples carrying these pathogenic variants, no variants showed haplogroup preference in this Chinese cohort (Figure S4E).

### 12 mtDNA variants significantly associated with nDNA variants

Genes encoded by mtDNA need to interact with some genes encoded by nDNA to exert functions, so there may be some extent of correlations between them. Studies have reported that the interaction between mtDNA and nDNA variations can affect the development and progression of complex diseases, such as cardiomyopathy, Parkinson’s disease, Alzheimer’s disease [58–61]. To explore the association of mtDNA-nDNA variants, we conducted genome-wide association analyses using 3945 unrelated samples from NyuWa, investigating 93 common mtDNA (minor allele frequency (MAF) ≥ 5%) and the genome-wide 7,124,343 nDNA variants (MAF ≥ 5%) (see detals in the Methods). After adjusting for multiple comparisons (FDR_BH < 0.05/93 = 5.4 × 10^−4^), we discovered a significant association between 12 mtDNA variants and 199 nDNA variants (Table S4). 9 of 12 mtDNA variants occurred in protein-coding genes, and they were all marker variants of haplogroups in Phylotree [62]. Of the 12 mtDNA variants significantly associated with nDNA variants, 5 variants were homoplasmic (Figure S7A). Among the 9 mtDNA variants in protein-coding genes, m.4769A>G located in gene *MT-ND2* had the highest population frequency in NyuWa (Figure S7B-C). And m.4769A>G exhibits the most significant association with nDNA variants (Figure 4A). Peifang Jiang et al. found that m.4769A>G is significantly associated with Tic disorders (TDs) [63]. Ludwig-Slomczynska AH et al. found that m.4769A>G interacted with seven nDNA variants (rs2255074, rs1107479, rs7973157, rs2950390, rs2958154, rs2290893, rs2035081) [64]. Besides, 199 nDNA variants are located within or in close proximity to 145 distinct genes. We performed pathway enrichment analysis, yet no significant pathway enrichment was observed. Genes located within a 25kb upstream and downstream range of the 199 nDNA variants were significantly enriched in pathways related to lipid metabolism and inflammatory response (Figure S7D). When applying a more stringent threshold (*P* = 5.0 × 10^-8^/93 = 5.4 × 10^-10^), we identified 82 significant associations between 8 mtDNA variants and 59 nDNA variants.

**Figure 4.**
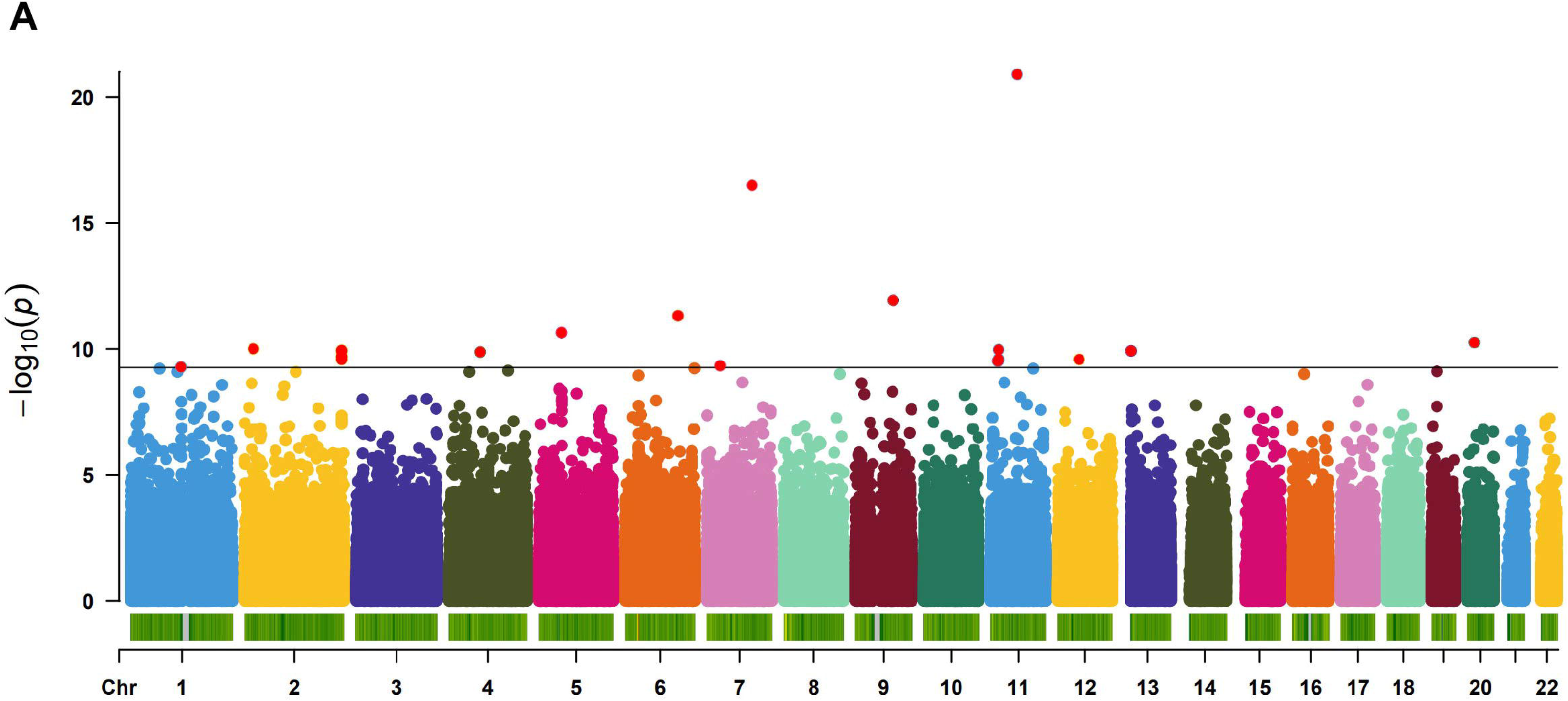
The mtDNA variant m.4769A>G significantly associated with nDNA variants. **A.** Manhattan plot of the association *P* values for mtDNA variant m.9950T>C and nDNA variants. The horizontal lines represent the study-wide significant threshold 5.4 × 10^-10^.

### Haplogroup composition of the Chinese population

Due to the maternal inheritance and lack of recombination of mtDNA, haplogroups categorize mtDNA haplotypes based on distinct sets of characteristic variants. Haplotype analysis is essential for mtDNA variant studies, providing critical insights into the genetic lineage and historical population migration [65]. Also, it’s crucial for correctly understanding the function of mtDNA variants, especially in disease and population genetics [66]. By constructing haplotypes, we can trace the inheritance patterns of mtDNA variants, identify Chinese population-specific frequency spectra, and reveal heterogeneity across different geographic regions in China. Based on Phylotree v17 [62], the most likely haplogroup of each individual was assigned using Haplogrep [67]. Among the 5184 haplogroups included in the Phylotree database [62], 1563 haplogroups were found in NyuWa and 1KGP (Figure S8A). 99.59% of our samples had quality score greater than 0.8 based on haplogrep results (Figure S8B). According to this, 7266 samples could be classified into 22 macrohaplogroups, and 72.53% (5,270/7,266) were assigned to Asia haplogroups based on the population divisions of MITOMAP (https://www.mitomap.org/foswiki/pub/MITOMAP/WebHome/simple-tree-mitomap-2019.pdf) (Figure 5A). Among the 22 macrohaplogroups, group D contained the largest proportion of samples, followed by M, B, and F of Asia haplogroups., As a descendant haplogroup of haplogroup M, haplogroup D is believed to have arisen in Central Asia (https://haplogroup.org/mtdna/rsrs/l123456/l23456/l2346/l346/l34/l3/m/m80d/d/). Moreover, there were 113 samples of NyuWa belonged as European haplogroups. Of the 113 samples, 78 were annotated to macrohaplogroup R, which is a child of haplogroup N. The haplogroup R originates in the West Asia [68], and dominates the European maternal landscape, making up 75-95% of the lineages there (https://haplogroup.org/mtdna/rsrs/l123456/l23456/l2346/l346/l34/l3/n/r/). Of our cohort, 5.11% of the samples belonged to macrohaplogroup H, which is more prevalent in Europe (Figure 5A).

**Figure 5.**
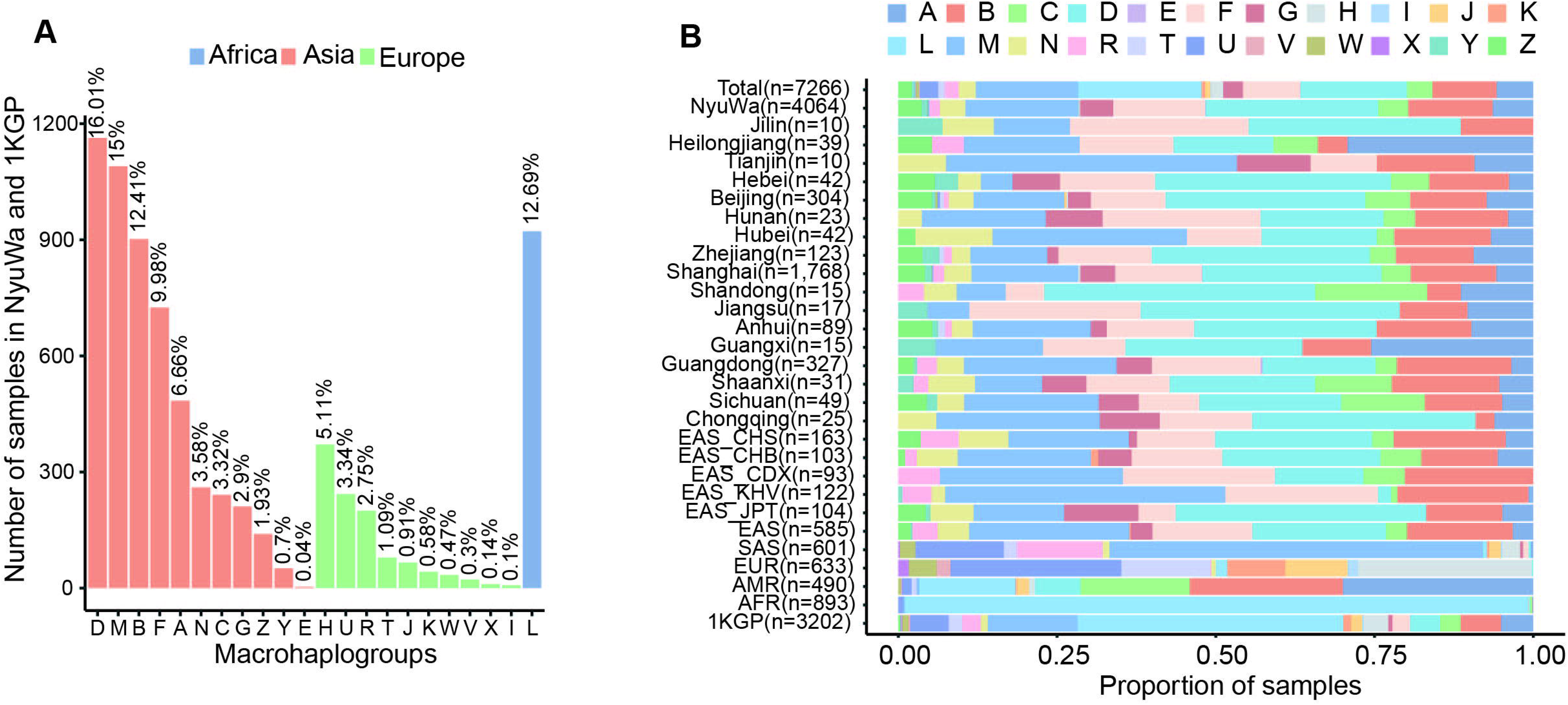
Haplogroup composition in this study. A. The number of samples under each macrohaplogroup in NyuWa and 1KGP. Colors indicated haplogroups associated with Asian and European, according to the population division of MITOMAP. B. The composition of macrohaplogroups in each administrative region.

The definition of Phylotree about haplogroup determines the difference in the number of mtDNA variants among different ethnic groups [37]. We analyzed the total number of mtDNA variants in each sample according to macrohaplogroup. In our cohorts, there were 20-50 mtDNA variants in all samples except macrohaplogroup H (Figure S8C), and the distributions of the number of variants were similar when focusing on homoplasmic mtDNA variants (Figure S8D). However, the numbers of heteroplasmic variants were similar across all the assigned macrohaplogroups (Figure S8E).

To obtain the composition of haplogroups in geographical regions in China, the proportion of each macrohaplogroup was calculated in administrative regions containing more than 10 samples (Figure S8F). Macrohaplogroups A, B, D, F, and M accounted for more than 3% frequency of the corresponding administrative regions (Figure 5B and S8G). The composition of NyuWa macrohaplogroups was similar to that in 1KGP East Asian (EAS) (Figure 5B). In Heilongjiang and Guangxi, macrohaplogroup A was more prevalent, similar to the proportion in American. However, this composition may be due to the limited sample size. Given that the NyuWa samples primarily originate from Beijing, Shanghai, and Guangdong, with each region contributing over 300 samples, the macrohaplogroup compositions of these regions exhibit similarities, yet frequency differences are also observed. We conducted an analysis of the mtDNA variation landscape across these three regions (Figure S9). The patterns of mtDNA variants and sample distribution in Beijing, Shanghai, and Guangdong showed similarities, which was consistent with the trend of the overall NyuWa (Figure S9A-F). Although differences in sample size may have caused some differences, these regions also have their own specific variants (Figure S9G).

### Relatively higher stability of rRNAs and tRNAs than other mtDNA regions

As previously described [38], we extracted regions with no mtDNA variants in this cohort, and explored whether there were specific or particularly conserved regions that remain invariant in the mtDNA across the Chinese population. Only considering SNVs and deletions, which could lead to the change of existing nucleotides of mtDNA sequence, 61.08% of mtDNA nucleotides were invariable and relatively stable in this cohort (Figure 6A). These stable nucleotides were more conserved compared with variable nucleotides (Figure S10A-B). rRNA and tRNA gene regions had a higher proportion of invariable nucleotides than protein-coding and non-gene regions (Figure 6B). Similar results were obtained when invariable nucleotides were extended to invariable intervals with insertions taken into account. There were 2503 invariable intervals (>1 nt) detected, 60 of which were longer than 10 nt (Figure S10C). In this cohort, rRNA and tRNA genes contained longer invariable intervals compared with others regions (Figure 6C). The longest invariable interval came from protein_coding gene, *MT-ND5*. To explore relatively stable segments shared across different populations, we performed an integrated analysis of invariable intervals in NyuWa, gnomAD, and HelixMTdb. As a result, we obtained 7 shared invariable intervals of length greater than 10 nt, which were locating in 2 rRNA genes, 4 tRNA genes, and 1 intergenic region (Table S5). Compared to other regions, the regions encoding rRNAs and tRNAs were more stable.

**Figure 6.**
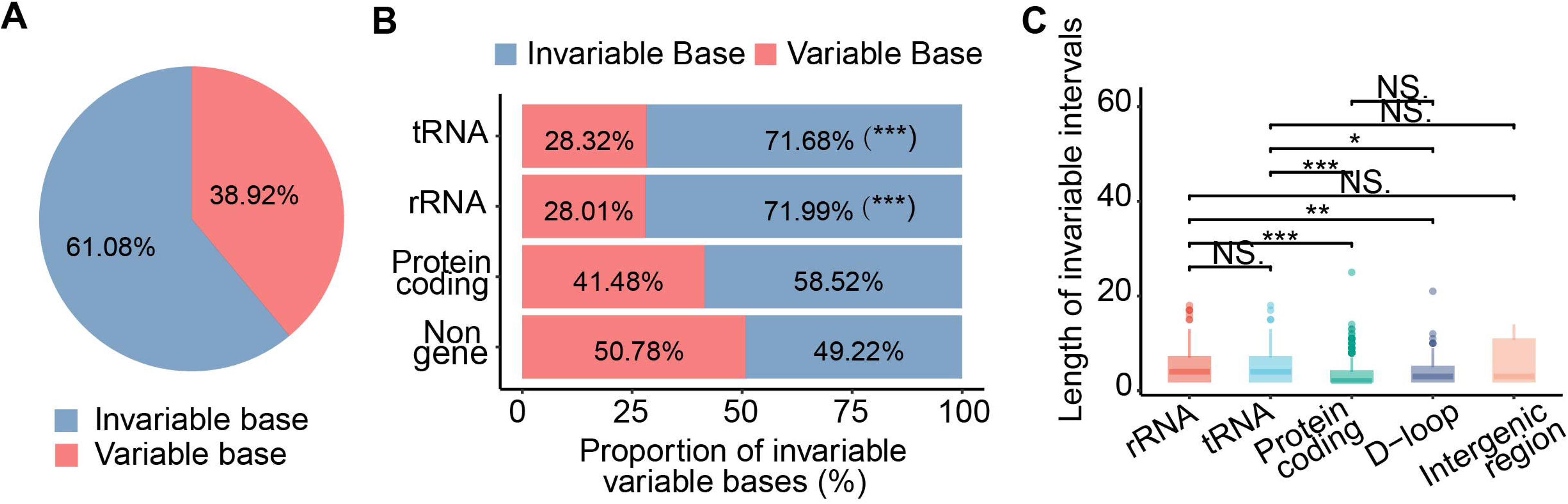
Invariable segments in mtDNA. A. Proportion of invariable and variable bases in mtDNA of this study. B. Proportion of invariable/variable bases in different mtDNA regions of this study. *** indicating *P* < 0.001 based on one-tailed Hypergeometric test. C. The length distribution in different mtDNA regions of this study. * for *P* <= 0.05, ** for *P* <= 0.01, *** for *P* <= 0.001, NS. for no significance based on one-tailed Wilcoxon test.

### Characteristics of NUMT insertions

NUMTs can be derived from any part of the mtDNA, while there is no consensus on the mechanism by which fragmented mtDNA exits mitochondria and inserts into the nuclear genome [69]. Here, we focused on the enrichment of NUMT breakpoints on mtDNA and nuclear genomes. Usually one NUMT corresponds to two mtDNA breakpoints. The filtered mtDNA breakpoints were likely to involve *ND5* and *RNR1* regions (Figure 7A). *RNR1* encode rRNA and the *RNR1* region is observed to have a higher normalized NUMT mtDNA breakpoints count compared to other regions of similar length. *ND5* is a mutational ‘hot spot’ and encode for the NADH-ubiquinone oxidoreductase chain 5 protein [70]. Regarding the *ND5* region, its longer mtDNA region length contributes to a greater number of NUMT mtDNA breakpoints. Breakpoints of common NUMTs were enriched in the coding regions (CDS) in the middle of the mtDNA, while rare NUMTs’ breakpoints tended to occur on both sides of the coordinate position including D-loop region (Figure 7B). Similarly, different categories of NUMTs have different characteristics at their insertion positions in the nuclear genome and NUMTs were found on all nuclear chromosomes (Figure 7C). Overall, the three datasets show similar trends in the enrichment patterns. Common NUMTs tended to insert in genomic duplications, simple repeats, miRNA and snoRNA. Breakpoints of rare NUMTs were accumulated in LINEs (Figure 7D). Considering that ultra-rare NUMTs accounted for the majority, the overall breakpoints enrichment were consistent with ultra-rare NUMT breakpoints, mainly concentrated in gene regions (Figure 7D). Focusing on the NyuWa dataset, particularly the rare category, there is a distinct and significant enrichment observed in the LINE genomic feature (*P* = 0.004) (Figure 7D).

**Figure 7.**
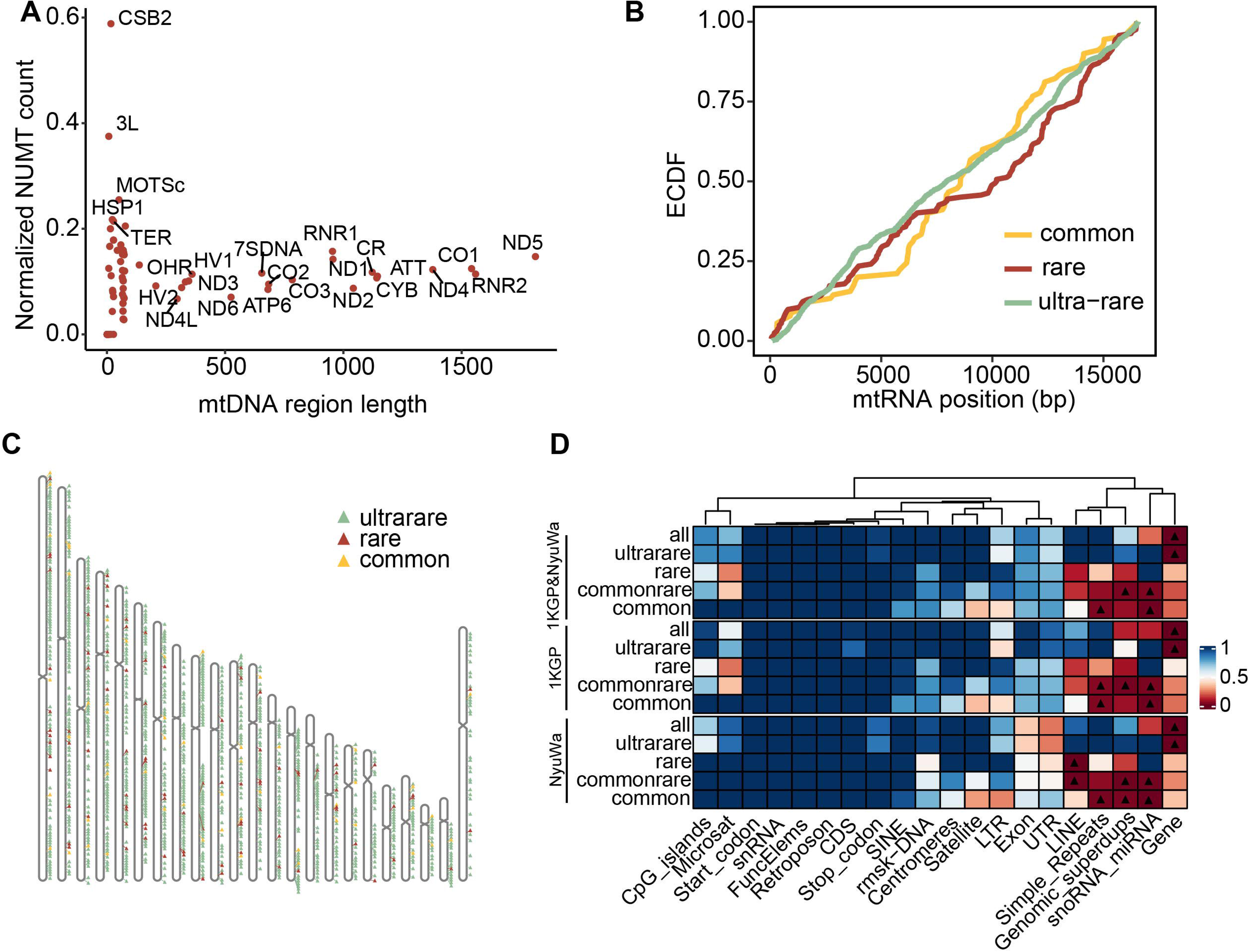
Characteristics of NUMT insertions. **A**. Comparison of the normalized number of NUMT mitochondrial breakpoints in mtDNA function locations. Normalized counts in regions with smaller intervals are greatly affected by randomness and are not considered here. **B.** ECDF plot of NUMT mitochondrial breakpoints. ECDF, Empirical cumulative distribution function. **C.** Chromosome map of chromosomal locations NUMTs inserted. **D.** P values for enrichment analysis of different genome regions. The “commonrare” refers to a combined group that includes both common and rare NUMTs. Triangle for *P* <= 0.05. snRNA, small nuclear RNA; Microsat, microsatellite; LTR, retrotransposon; rmsk-DNA, repetitive DNA; superdups, superduplications, LINE, long interspersed nuclear element; UTR, untranslated region.

## Discussion

WGS data can be used not only for the analysis of nuclear genome variation, but also for the discovery of mtDNA variation and NUMTs [23,71,72]. In addition, increasing studies have indicated that mtDNA variants are closely related to a wide variety of diseases [73–75]. Moreover, as an ongoing phenomenon, the integration of NUMTs mostly manifests as neutral polymorphism [76], while NUMTs were also associated with human diseases [77], various cancers [22], human lifespan [24] and introduce false positives variants in mitochondrial genetic variant datasets [58]. Here, we not only conducted a comprehensive analysis of mtDNA variation using deep WGS data from 7266 individuals, including 4064 samples from NyuWa Project [42] and 3202 samples from 1KGP, but also profiled the landscape of the NUMTs using 7324 samples, including 4122 samples from NyuWa Project [42] and 3202 samples from 1KGP. We detected 7216 distinct high-quality mtDNA variants with VAF >= 0.1 and found that the vast majority of variants were homoplasmic variants, which was consistent with the result of previous study [37]. We found 88 NyuWa-specific mtDNA variants by comparing with other mtDNA variants resources. The researches have indicated that mtDNA is associated with nDNA in human diseases and mtDNA genotypes can have effect on nDNA mutations [58]. We performed a genome-wide association analysis between mtDNA variants and nDNA variants to explore the association of mtDNA-nDNA in Chinese population. In NyuWa mtDNA resource, 12 mtDNA variants were significantly associated with 199 nDNA variants. Given the reported interactions between mtDNA and nDNA, we may provide a reference for the association of mtDNA-nDNA variants. Besides further investigation would be necessary to elucidate their associations. However, in the previous association analysis of mtDNA-nDNA in Japanese population, no significant association was found between mtDNA and nDNA [39]. This may be due to the limited number of samples. There are 1928 individuals in Japanese cohort while 4064 in NyuWa. On the other hand, it may also be related to the stratification of the population.

m.4769A>G exhibits the most significant association with nDNA variants. Peifang Jiang et al. found that m.4769A>G is significantly associated with TDs based on an additive model [63]. Besides, among the 12 mtDNA variants obtained by mtDNA-nDNA association analysis in NyuWa, m.14766C>T, m.7028C>T, and m.750A>G were also reported to be associated with TDs by Peifang Jiang et al. A. Jeremy Willsey et al. reported nDNA variants significantly associated with TD [78]. The nuclear genes where these variants are located overlap with the genes where the nuclear SNPs significantly associated with m.4769A>G, m.14766C>T, m.7028C>T, and m.750A>G are located, such as *DYNC2H1, HHIPL2, KIAA1549L, NAV2* and *SLC1A3*. After annotation, 199 nDNA variants are located within or in close proximity to 145 distinct genes. Upon intersecting these 145 genes with 2,282 MT-nDNA candidate genes (Aldi T Kraja et.al., Am J Hum Genet., 2018) [79], we identified a subset of 10 mitochondrial-associated nuclear genes: *ATG4D, MIMT1, NAV2, NUCB2, PECAM1, CALR, ECHDC3, FKBP10, STARD7*, and *TAT*. These 10 mitochondrial-related nuclear genes play an important role in cellular energy metabolism, autophagy, and mitochondrial homeostasis. Then we performed pathway enrichment analysis on 145 distinct genes related to 199 nDNA variants, yet no significant pathway enrichment was observed. Genes located within a 25kb upstream and downstream range of the 199 nDNA variants were significantly enriched in pathways related to lipid metabolism and inflammatory response. This suggests that these genes may play an important role in regulating lipid levels, cholesterol storage, and the biosynthesis of inflammatory mediators. It has been previously suggested that mtDNA mutations are closely tied to lipid homeostasis [80], cholesterol metabolism [81] and inflammation [82]. These mutations can potentially lead to metabolic disorders [83]. Further study of the mechanism and function of the association of mtDNA-nDNA variants will help to gain a deeper understanding of the pathogenesis of mitochondrial-related diseases and provide new ideas for the diagnosis and treatment of related diseases.

For haplogroup analysis, we identified 1563 distinct haplogroups among the 5184 haplogroups included in the Phylotree database and classify 7266 samples into 22 macrohaplogroups, with 72.53% (5270/7266) assigned to Asian haplogroups. The most prevalent macrohaplogroups were D, M, B, and F, reflecting the significant genetic diversity within the Asian population. Notably, haplogroup D, a descendant of haplogroup M believed to have originated in Central Asia, was the most common. The geographic distribution of haplogroups within China revealed distinct patterns. Macrohaplogroups A, B, D, F, and M accounted for more than 3% of the samples in several administrative regions and showed the landscape of mtDNA variants in Beijing, Shanghai and Guangdong. The composition of haplogroups in some administrative regions is influenced by the limited sample size. Our study highlights the rich genetic diversity of mtDNA haplogroups in the Chinese population and provides a view of their geographic distribution and ancestral origins. In addition to the analysis of mtDNA variants, we also selected Beijing, Shanghai, and Guangdong as representative regions to illustrate the landscape of NUMTs across different administrative regions. This analysis is presented in Figure S11. The NUMT landscapes of Beijing, Shanghai, and Guangdong display similarities, which are consistent with the overall trends observed in the entire NyuWa dataset. Moreover, each region also harbors specific NUMTs.

It has been found that heteroplasmies throughout the mtDNA are common innormal human cells, and moreover, the frequencies of the heteroplasmic variants varied considerably between different tissues in the same individual [84]. mtDNA are extracted from blood samples in NyuWa [42], and tissue samples heavily enriched with mitochondria are not included. The mtDNA variant spectrum of this Chinese cohort can only reflect the heterogeneity level of mtDNA in blood.

For the NUMTs, there were a total of 120,400 NUMTs in the 1KGP and NyuWa and 1466 distinct NUMTs were identified after deduplication between 7324 samples. The length of NUMTs inserted into the nuclear genome ranged from 5bp to the entire mtDNA, and we didn’t consider concatenated NUMTs and other complex situations. We show that NUMTs exhibited diversity among the NyuWa Chinese population, with ultrarare NUMTs accounting for 85.87% of the total, but only appearing in 19.36% of the samples. The NyuWa and 1KGP show similar trends in the nuclear breakpoint enrichment patterns. Overall, the breakpoint distribution of ultrarare NUMTs on the mtDNA was relatively random, and they mainly inserted into the gene regions of the nuclear genome, which may explain the low population frequency. Meanwhile, mtDNA breakpoints of common and rare NUMTs exhibited clear enrichment tendency. Common NUMTs had a relatively high cumulative distribution slope in the middle mtDNA region, while rare NUMTs breakpoints tended to occur on both sides including D-loop region. Considering the high population frequency, common NUMTs nuclear breakpoints accumulated in repetitive regions. Besides, rare NUMTs tend to insert into LINE regions. Focusing on the NyuWa dataset, particularly the rare category, there is a distinct and significant enrichment observed in the LINE genomic feature (*P* = 0.004). There are some divergences from the published results of 100,000 Genomes Project (mostly European), such as the abundance of regulatory elements, short interspersed elements (SINEs), simple repeats, and introns in ultrarare NUMTs’ nuclear breakpoints [23]. Moreover, the comparison of normalized breakpoints counts in various functional regions of mtDNA requires consideration of interval length, as normalized count in regions with small intervals is greatly affected by randomness.

However, the uneven distribution of samples across various geographical regions within the NyuWa cohort constrains our comprehensive analysis of haplogroup composition in these areas. Furthermore, the cohort’s representative power could be significantly enhanced by augmenting the sample size from these underrepresented areas.

In general, we provide valuable reference resources for genetic researches on mtDNA-related diseases, especially East Asian populations. Our previous work, which established a comprehensive WGS resource for the Chinese population, has already laid a solid foundation for this endeavor. In the NyuWa project, we meticulously described the nuclear SNPs and indels [42], mobile element insertions (MEIs) [43], and short tandem repeats (STRs) [44]. Looking ahead, with the integration of these data, we will be better equipped to dissect the complex interplay between nuclear and mitochondrial genomes, associations between complex variations and their implications in disease susceptibility and progression.

## Materials and methods

### Samples and mtDNA variants calling

This study was approved by the Medical Research Ethics Committee of Institute of Biophysics, Chinese Academy of Sciences and complies with all relevant ethical regulations. All participants provided written informed consent. The informed consent is used to collect samples for genome studies conducted by Chinese Academy of Sciences. 7331 samples in this study were collected from the NyuWa Project and the 1KGP. Of these, 4129 samples of the NyuWa [42] were from different administrative regions of China. WGS was performed using Illumina platform and the sequencing data were aligned to GRCh38 (containing mtDNA, NC_012920.1) to get bam files using BWA-mem v0.7.15 [85], which was also used in the GATK Best Practices Workflows [86,87]. We used mitochondrial variants calling pipeline (https://github.com/gatk-workflows/gatk4-mitochondria-pipeline) with the default parameters to identify mtDNA variants for bam file of each sample [37]. The mtDNA variant calling was performed on 1KGP using the same workflow as above.

### Acquisition of high-quality mtDNA variant set

To obtain high-quality mtDNA variants, the samples and variants were filtered separately. Sample filtration was performed as follows. 1) Using VerifyBamID2 [88] version 1.0.6 to assess contamination based on nDNA, samples were excluded with contamination > 2%. 2) Samples reported as contaminated by haplocheck v1.3.2, a tool to identify contamination using the mitochondrial phylogeny [45], were removed. 3) Samples with mtDNA copy number < 50 were removed [37]. The mtDNA copy number for each sample was calculated with the following formula: mtDNA copy number = (Mean mtDNA depth/Mean autosomal depth) * 2 [39]. 4) The duplicate data, possibly from the same persons, were discarded according to the method described in the previous article [42]. Ultimately, Samples used for calling mtDNA variants after filtering included 4064 Chinese samples and 3202 1KGP. For mtDNA variants filtering, we excluded variants in known WGS artifacts and low-complexity sites (66-71, 301, 302, 310, 316, 3107, 12418-12425, 16182-16194). The mtDNA variants marked as ‘PASS’ in the final VCF file generated by mitochondrial variants calling pipeline and with the depth greater than 100X, were selected. On this basis, the filtered variants with VAF (ratio of reads supporting variant to all reads aligned to this locus) >= 0.1 were assigned as high-quality variants in this cohort. A total of 268,746 high-quality variants at different heteroplasmic levels were obtained. After deduplication, the final distinct high-quality mtDNA variant set contained 7216 variants with VAF >= 0.1 from 4064 NyuWa samples and 3202 1KGP samples.

### Characteristic analysis of mtDNA variants

VAF is calculated by dividing the Allele Depth (AD), which represents the number of sequencing reads supporting the variant allele in the genome, by the total sequencing depth (DP), which represents the total number of reads covering the variant site. Based on VAF, the high-quality mtDNA variant set was categorized into two groups: homoplasmic variants (VAF ranging from 0.95 to 1.00) and heteroplasmic variants (VAF ranging from 0.10 to 0.95) [37]. Homoplasmic variants indicate that the variant is present in nearly all mtDNA copies within the individual, while heteroplasmic variants reflect the coexistence of both variant and wild-type mtDNA copies. Because a mtDNA variant may have different VAFs in different individuals, the distinct high-quality mtDNA variants were divided into three categories: detected only at homoplasmic level, detected only at heteroplasmic level, detected both at homoplasmic and heteroplasmic levels (defined as both in this study). In addition to classifying variants, we also analyzed the frequency of these mtDNA variants in the population. The population-level allele frequency reflects the prevalence of a variant across multiple individuals. Based on these analyses, we described the characteristics of the variants. Circos [89] v0.69.5 was used to visualize the variants of mtDNA resource in mtDNA.

### Functional annotation of mtDNA variants

The annotation of mtDNA variants was performed using Variant Effect Predictor (VEP v101) [53], with parameters ‘--everything --flag_pick_allele --distance 0’. For the variants in protein-coding genes, we screened for missense mutations among the 7216 high-quality mtDNA variant sites and performed pathogenicity annotation based on the APOGEE2 predictor results from the MitImpact database (Version 3.1.2) [54]. The pathogenicity of the tRNA variants was classified as ‘Likely Pathogenic’ with >16.25 (75%-100%), ‘Possibly Pathogenic’ with 12.66-16.25 (50%-75%), ‘Possibly Benign’ with 8.44-12.66 (25%-50%), and ‘Likely Benign’ with <8.44 (0%-25%) by MitoTip [57] score, which was obtained by extracting and submitting the related tRNA variants to the Mitomaster Web API (https://www.mitomap.org/foswiki/bin/view/MITOMASTER/WebHome) [38].

### Comparison with other mtDNA variant resources

We downloaded the mtDNA variants from gnomAD on January 17, 2022 (https://gnomad.broadinstitute.org/downloads#v3-mitochondrial-dna), HelixMTdb (https://helix-research-public.s3.amazonaws.com/mito/HelixMTdb_20200327.tsv), MITOMAP on December 28, 2023 (https://www.mitomap.org/MITOMAP). After dividing multi-allele and removing duplicate variants, the final number of mtDNA variants was 10,850 from gnomAD, 11,491 from HelixMTdb, and 19,092 from MITOMAP. We compared the high-quality mtDNA variants of NyuWa mtDNA resource with other four resources in terms of shared variants, SNV types, and the correlation of variant frequency in populations.

### Analysis of mtDNA-nDNA variant association

To analyze the association between mtDNA variants and nDNA variants, we eliminated samples of possible relatives as the method described in NyuWa Genome resource [42]. 3,945 unrelated samples were used for the association analysis using mtDNA variant heteroplasmy levels as a continuous phenotype. 93 mtDNA variants with major allele frequency >5% were chosen from the unrelated samples, as well as 7,090,977 nDNA variants with sample call rate > 0.9, variant call rate > 0.9, MAF > 5%, and Hardy-Weinberg equilibrium P >= 1.0 × 10^-6^ were extracted from the same samples. The analysis of mtDNA-nDNA variants association was conducted using glm by PLINK v2.00a5.10 with top 12 principal components (PCs) of nDNA variants and sex as covariables. The sex of all samples was inferred based on chrX Ploidy and chromosome depth. Method details for sex inference described in NyuWa Genome resource [42]. Considering the multiple comparison of the mtDNA variants, FDR_BH < 0.05/93 = 5.4 × 10^−4^ and *P* < 5 × 10^-8^/93 = 5.4 × 10^−10^ were regarded as the threshold for significant association [39].

### Annotation of mtDNA haplogroups

Haplogrep [67] was used to assign haplogroup to each sample based on Phylotree v17 [62]. To meet the right-alignment of indels in Phylotree, left-alignment mtDNA indels in our cohort were manually changed to right-alignment. We run haplogrep using default parameters except parameters ‘--hetLevel 0.1 --extend-report’. Only the first ranked haplogroup of each sample was reported and used for subsequent analysis. To obtain the composition of mtDNA haplogroups in different administrative regions of China, the macrohaplogroups (first letter of the haplogroup reported by haplogrep) of each individual were extracted and statistically analyzed, and compared with 1KGP haplogroups which defined with the same method.

### Pathogenic mtDNA variants in NyuWa mtDNA resource

The pathogenic mtDNA variants were downloaded from MITOMAP [90] on July 27, 2021, and the variants labeled as ‘Confirmed’ were picked. 96 confirmed pathogenic variants were used for query existence, frequency, and heteroplasmic levels in NyuWa and 1KGP samples.

### Stability Analysis in mtDNA regions

To analyze the invariable bases in NyuWa mtDNA resource, the variable bases detected were excluded from mtDNA sequence. The detailed method was that only SNVs and deletions, which could change the existing mtDNA bases, were considered. We deduplicated the variable bases and extracted the invariable bases from mtDNA sequence in NyuWa mtDNA resource. The invariable and variable bases were annotated to mtDNA regions using BEDTools intersect [91]. The PhastCons scores and phyloP scores, which were downloaded from UCSC [92], were used to evaluate the conservativeness of invariable bases. The invariable bases were expanded into intervals by in house script, and insertions were also considered. Intervals without variation were defined as invariable intervals in NyuWa mtDNA resource. In order to obtain NyuWa-specific invariable intervals, the comparative analysis was performed among NyuWa and gnomAD, HelixMTdb.

### Detecting NUMTs

NUMTs were not detected in gametes, but presumably in blood samples. On the basis of removing contaminated samples and duplicate data, we removed samples according to insert size (<250 bp). After quality control, 7324 samples’ WGS bam files were processed to generate discordant reads and split reads using samblaster. We treated discordant reads within 500bp as a cluster. The cluster supported by less than five pairs of discordant reads are discarded and the NUMTs within a distance of 1,000 bp from multiple samples were grouped as the same NUMT. Then the split reads within 500 bp NUMT flanks (upstream 500 bp + downstream 500 bp) of each cluster were realigned to the human reference genome (hg38) using BLAT. The output of blat were used to search mtDNA and nuclear breakpoints which were determined by filtering at least two split reads. Here we didn’t consider concatenated NUMTs and other complex situation.

Bedtools was used to detect known NUMTs, which were downloaded from UCSC and previous publications [23,49–52]. 500 base pairs added upstream and downstream of each NUMT on the nuclear genome in the published NUMT datasets when searching for overlaps in NyuWa NUMT dataset.

All steps above are performed using the code from the github public NUMT pipeline (https://github.com/WeiWei060512/NUMTs-detection.git) described in the NUMTs of the 100,000 Genomes Project study (Wei Wei et al., 2022) for our Nyuwa genome dataset.

### mtDNA and nuclear breakpoints enrichment analysis

For distribution of NUMTs breakpoints on mtDNA, the cumulative distribution was implemented to observe the slope of NUMT breakpoints at different positions of mtDNA, reflecting the degree of breakpoints enrichment.

To analysis NUMT insertion nuclear sequence characteristics, genome regions (centromere, genomic duplications, simple repeats, functional elements, CpG islands, satellites, LINEs, SINEs, gene, etc) were downloaded from UCSC (https://genome.ucsc.edu/) and also extracted from the gencode annotation gtf file (version43). Resampling 10,000 sets of random nuclear positions in 200 bp NUMT flanks (upstream 100 bp + downstream 100 bp) based on total number of NUMTs (permutation test) was performed to generate empirical P values by comparing with the observed NUMTs in target regions.

### Database development

NMVR was developed by Bootstrap and PHP. Data resources were stored in the MySQL database. Users can browse the information of NMVR at http://bigdata.ibp.ac.cn/NMVR/.

### Quantification and statistical analysis

All statistical analyses in this study were performed using R v4.1.0 and the statistical tests were described in figure legends and the Methods section. Unless otherwise indicated, statistical significance testing was one-sided.

## Data and code availability

The DNA sequencing data of NyuWa samples used in this study have been deposited in the Genome Sequence Archive (GSA) in National Genomics Data Center, China National Center for Bioinformation/Beijing Institute of Genomics, Chinese Academy of Sciences, under accession number HRA004185 (https://ngdc.cncb.ac.cn/gsa-human/). These data are available under restricted access for privacy protection and can be obtained by application on the GSA database website (https://ngdc.cncb.ac.cn/gsa-human/) following the guidance of “Request Data” on this website. These data have also been deposited in the National Omics Data Encyclopedia (NODE) of the Bio-Med Big Data Center, Shanghai Institute of Nutrition and Health, Chinese Academy of Sciences, under accession number OEP002803 (http://www.biosino.org/node). The user can register and login to this website and follow the guidance of “Request for Restricted Data” to request the data. The alignment files for the 1KGP dataset are available at https://ftp-trace.ncbi.nlm.nih.gov/1000genomes/ftp/1000G_2504_high_coverage/data/ and https://ftp-trace.ncbi.nlm.nih.gov/1000genomes/ftp/1000G_2504_high_coverage/addit ional_698-_related/. Codes used in the study are available at https://github.com/gatk-workflows/gatk4-mitochondria-pipeline and https://github.com/WeiWei060512/NUMTs-detection.git.

## CRediT author statement

**Yuanxin Wang:** Conceptualization, Investigation, Methodology, Formal analysis, Visualization, Writing – original draft, Writing – review & editing. **Jiajia Wang:** Conceptualization, Investigation, Methodology, Formal analysis, Visualization, Writing – original draft, Writing – review & editing. **Yanyan Li:** Data curation, Methodology, Formal analysis. **Peng Zhang:** Data curation, Investigation, Methodology. **Zhonglong Wang:** Formal analysis, Visualization. **Shuai Liu:** Data curation. **Yiwei Niu:** Data curation. **Yirong Shi:** Data curation. **Sijia Zhang:** Data curation. **Tingrui Song:** Data curation. **Tao Xu:** Conceptualization, Funding acquisition, Project administration, Resources. **Shunmin He:** Conceptualization, Funding acquisition, Project administration, Resources, Writing – review & editing. All authors read and approved the final manuscript.

## Acknowledgements

We thank Center for Big Data Research in Health (http://bigdata.ibp.ac.cn/), Institute of Biophysics, Chinese Academy of Sciences, for supporting data analysis and computing resource. This work was supported by Beijing Natural Science Foundation [L248016 (W.-M.Z.)]; National Key R&D Program of China [2021YFF0704500 (P.Z.), 2022YFC3400405 (S.-M.H.)]; 14th Five-year Informatization Plan of Chinese Academy of Sciences [CAS-WX2021SF-0203 (S.-M.H.)]; National Natural Science Foundation of China [91940306 (S.-M.H.), 31970647 (P.Z.), 32200478 (Y.-Y.L.)] and China Postdoctoral Science Foundation [2022M713311 (Y.-Y.L.), GZC20232899 (G.-Y.W.)].

## Supplementary material

**Figure S1 High-quality mtDNA variants**

**A.** Distribution of nDNA mean depth in individuals (n = 7331). **B.** Distribution of mtDNA mean depth in individuals(n = 7331). **C**. Distribution of copy number of mtDNA in this cohort. **D.** Correlation of autosomal mean depth and mtDNA mean depth. **E.** The ratio of nDNA to mtDNA depth in all samples of this cohort. **F.** Distribution of high-quality mtDNA variants for each sample in this study.

**Figure S2 Characteristics of mtDNA variants**

**A.** Proportion of homoplasmic and heteroplasmic variants in total mtDNA variants. **B.** Proportion of the transition and transversion SNVs in this study. **C.** Proportion heatmap of transition and transversion SNVs. The horizontal axis represents alt bases, and the vertical axis represents ref bases. The denominators of each row of the matrix are the total number of A, T, C, and G on the mitochondrial reference genome. **D.** Length distribution of mtDNA indels in this study. **E.** Variant types of non-redundant high-quality mtDNA variants detected at different variant frequencies in this study. **F.** Variant types of non-redundant high-quality mtDNA variants detected at different heteroplamic levels (homoplasmic, heteroplasmic, and both). **G**. Distribution of variants only observed at heteroplasmic levels in this study.

**Figure S3 Comparison of mtDNA variants in NyuWa with other resources**

**A.** Frequency of variants between NyuWa and 1KGP. Pearson correlation coefficients were shown in the figure. **B**. Frequency of variants between NyuWa and HelixMTdb. **C**. Frequency of variants between NyuWa and gnomAD. **D.** Frequency of variants between NyuWa and AFR of 1KGP. AFR, African. **E.** Frequency of variants between NyuWa and AMR of 1KGP. AMR, American. **F.** Frequency of variants between NyuWa and EUR of 1KGP. EUR, European. **G.** Frequency of SNVs between NyuWa and gnomAD. **H.** Frequency of insertions between NyuWa and gnomAD. **I.** Frequency of deletions between NyuWa and gnomAD. **J.** Number of NyuWa-specific variants at different heteroplasmic levels in mtDNA regions. **K.** Shown all heteroplasmic levels of 88 NyuWa-specific variants in this cohort. Different colors represent different variants.

**Figure S4 Predicted functions and pathogenicity of mtDNA variants**

**A.** Number of indels in mtDNA regions. **B.** Number of mtDNA heteroplasmic variants in mtDNA regions. **C.** Proportion of variants (homoplasmic, heteroplasmic, and both) per protein-coding gene. **D.** Proportion of variants (homoplasmic, heteroplasmic, and both) per tRNA gene. **E.** Known pathogenic variants observed in this study along with the haplogroups.

**Figure S5 Predicted functions and pathogenicity of mtDNA variants in NyuWa**

**A.** Number of mtDNA variants annotated to the mtDNA regions. **B.** Proportion of variant annotations in mtDNA regions. mtDNA variants for total variants. Homoplasmic for variants observed only at homoplasmic levels. Heteroplasmic for variants observed only at heteroplasmic levels. Both for variants observed both at homoplasmic and heteroplasmic levels. Background for the mtDNA genome. ** for *P* <= 0.01, *** for *P* <= 0.001 based on one-tailed Hypergeometric test. **C.** The distribution of the maximum heteroplasmic levels of mtDNA variants in mtDNA regions. *** for *P* <= 0.001, ** for *P* <= 0.01, NS. for no significance based on one-tailed Wilcoxon test. **D.** Number of missense variants in protein-coding genes. APOGEE 2 refers to five pathogenicity classes: benign, likely-benign, VUS, likely-pathogenic, and pathogenic. VUS+ means closer to the likely pathogenic threshold. VUS-means closer to the likely benign threshold. **E.** Number of variants in tRNA genes. The severity of variants was defined by MitoTIP. Likely pathogenic: Variants with high pathogenicity scores, indicating a strong likelihood of causing disease. Possible pathogenic: Variants with moderate pathogenicity scores that may be associated with disease, suggesting a potential but less certain disease-causing effect. Possibly benign & likely benign: Variants with low pathogenicity scores, unlikely to cause disease. Unknown: Variants that do not have a MitoTIP percentile score.

**Figure S6 Predicted functions and pathogenicity of mtDNA variants in 1KGP**

**A.** Number of mtDNA variants annotated to the mtDNA regions. **B.** Proportion of variant annotations in mtDNA regions. mtDNA variants for total variants. Homoplasmic for variants observed only at homoplasmic levels. Heteroplasmic for variants observed only at heteroplasmic levels. Both for variants observed both at homoplasmic and heteroplasmic levels. Background for the mtDNA genome. ** for *P* <= 0.01, *** for *P* <= 0.001 based on one-tailed Hypergeometric test. **C.** The distribution of the maximum heteroplasmic levels of mtDNA variants in mtDNA regions. *** for *P* <= 0.001, ** for *P* <= 0.01, NS. for no significance based on one-tailed Wilcoxon test. **D.** Number of missense variants in protein-coding genes. APOGEE 2 refers to five pathogenicity classes: benign, likely-benign, VUS, likely-pathogenic, and pathogenic. VUS+ means closer to the likely pathogenic threshold. VUS-means closer to the likely benign threshold. **E.** Number of variants in tRNA genes. The severity of variants was defined by MitoTIP. Likely pathogenic: Variants with high pathogenicity scores, indicating a strong likelihood of causing disease. Possible pathogenic: Variants with moderate pathogenicity scores that may be associated with disease, suggesting a potential but less certain disease-causing effect. Possibly benign & likely benign: Variants with low pathogenicity scores, unlikely to cause disease. Unknown: Variants that do not have a MitoTIP percentile score.

**Figure S7 mtDNA variants significantly associated with nDNA variants**

**A.** Heteroplasmic levels of 12 mtDNA variants associated with nDNA variants. **B**. Number of samples containing the 12 mtDNA variants in this study. **C**. Population/Variant frequencies of mtDNA variant, m.4769A>G, in NyuWa, gnomAD, and 1KGP. The term “Total” refers to the 1KGP or genomAD entire dataset. The category ‘Other’ encompasses samples that do not belong to the listed populations (European, East Asian, Ashkenazi Jewish, American, and African) and includes additional populations not specifically detailed in the figure. D. Enriched gene ontology (GO) pathways for the genes located within a 25kb upstream and downstream range of the 199 nDNA variants.

**Figure S8 Haplogroup composition in this study (related to Figure 5**)

**A.** Venn diagram of overlapping mtDNA haplogroups in this study and Phylotree. **B.** The distribution of the quality scores for haplogroups based on Haplogrep software. Median and mean quality scores were shown in figure. **C.** Number of mtDNA variants across samples within each macrohaplogroup. Colors indicated haplogroups associated with Africa, Asian and European, according to the population division of MITOMAP. **D.** The number of homoplasmic variants across samples within each macrohaplogroup. Colors indicated haplogroups associated with Africa, Asian and European, according to the population division of MITOMAP. **E.** The number of heteroplamsic variants across samples within each macrohaplogroup. **F.** The number of samples in each administrative region in NyuWa. **G.** The composition of sample origins for each macrohaplogroup.

**Figure S9 Characteristics of mtDNA variants in Beijing, Shanghai and Guangdong**

**A.** Variant types (SNVs, deletions, insertions) of mtDNA variants detected at different administrative region (Beijing, Shanghai and Guangdong). **B.** Proportion of the transition and transversion SNVs in Beijing, Shanghai and Guangdong. **C.** Number of variants (homoplasmic, heteroplasmic, and both) in Beijing, Shanghai and Guangdong. Considering that Beijing and Guangdong only have about 300 samples, the heteroplasmic level classification of the variants were not re-performed here. **D.** Proportion of variants (homoplasmic, heteroplasmic, and both) in Beijing, Shanghai and Guangdong. **E.** Distribution of mtDNA variants per sample in Beijing, Shanghai and Guangdong. **F.** Number of heteroplasmic variants per sample in Beijing, Shanghai and Guangdong. **G.** Venn diagram of mtDNA variants distribution among Beijing, Shanghai, and Guangdong.

**Figure S10 Invariable segments in mtDNA**

**A.** phastCons scores of invariable and variable bases in mtDNA of this study. *** indicating *P* < 0.001 based on one-tailed Wilcoxon test. **B.** phyloP scores of invariable and variable bases in mtDNA of this study. *** indicating *P* < 0.001 based on one-tailed Wilcoxon test. **C.** The length distribution of the invariable intervals with length > 1 nt.

**Figure S11 Characteristics of NUMTs in Beijing, Shanghai and Guangdong**

**A.** The number distribution of NUMTs in Beijing, Shanghai and Guangdong. **B.** Size distribution of NUMTs less than 400 bp in Beijing, Shanghai and Guangdong. **C.** Proportion of samples carrying NUMTs by population frequency. Considering that Beijing and Guangdong only have about 300 samples, population frequencies of NUMTs were not recalculated here. **D.** Proportion of reported (known) and newly (unknown) identified NUMTs in Beijing, Shanghai and Guangdong. **E.** ECDF plot of NUMT mitochondrial breakpoints in Beijing, Shanghai and Guangdong. **F.** Chromosome map of chromosomal locations NUMTs inserted. **G.** Venn diagram of NUMTs distribution among Beijing, Shanghai, and Guangdong.

**Table S1 Information of 7331 samples in NyuWa and 1KGP**

**Table S2 Catalog of 13 pathogenic missense mutations**

**Table S3 Known pathogenic mtDNA variants present in 39 individuals**

**Table S4 Results of mtDNA-nDNA association analysis**

**Table S5 Shared invariable intervals within NyuWa, gnomAD, and HelixMTdb**

## References

[1] Taanman JW. The mitochondrial genome: structure, transcription, translation and replication. Biochim Biophys Acta 1999;1410:103–23.

[2] Healy TM, Burton RS. Strong selective effects of mitochondrial DNA on the nuclear genome. Proc Natl Acad Sci U S A 2020;117:6616–21.

[3] Anderson S, Bankier AT, Barrell BG, de Bruijn MH, Coulson AR, Drouin J, et al. Sequence and organization of the human mitochondrial genome. Nature 1981;290:457–65.

[4] Mishra P, Chan DC. Mitochondrial dynamics and inheritance during cell division, development and disease. Nat Rev Mol Cell Biol 2014;15:634–46.

[5] Clay Montier LL, Deng JJ, Bai Y. Number matters: control of mammalian mitochondrial DNA copy number. J Genet Genomics 2009;36:125–31.

[6] Wallace DC. Mitochondrial DNA variation in human radiation and disease. Cell 2015;163:33–8.

[7] Jr CWB. The Inheritance of Genes in Mitochondria and Chloroplasts: Laws, Mechanisms, and Models. Annu Rev Genet 2001;35:125–48.

[8] Xu S, Schaack S, Seyfert A, Choi E, Lynch M, Cristescu ME. High mutation rates in the mitochondrial genomes of Daphnia pulex. Mol Biol Evol 2012;29:763–9.

[9] Zaidi AA, Makova KD. Investigating mitonuclear interactions in human admixed populations. Nat Ecol Evol 2019;3:213–22.

[10] van den Ameele J, Li AYZ, Ma H, Chinnery PF. Mitochondrial heteroplasmy beyond the oocyte bottleneck. Semin Cell Dev Biol 2020;97:156–66.

[11] Stewart JB, Chinnery PF. The dynamics of mitochondrial DNA heteroplasmy: implications for human health and disease. Nat Rev Genet 2015;16:530–42.

[12] Picard M, Zhang J, Hancock S, Derbeneva O, Golhar R, Golik P, et al. Progressive increase in mtDNA 3243A>G heteroplasmy causes abrupt transcriptional reprogramming. Proc Natl Acad Sci U S A 2014;111:E4033–4042.

[13] Rossignol R, Faustin B, Rocher C, Malgat M, Mazat J-P, Letellier T. Mitochondrial threshold effects. Biochem J 2003;370:751–62.

[14] Craven L, Alston CL, Taylor RW, Turnbull DM. Recent Advances in Mitochondrial Disease. Annu Rev Genomics Hum Genet 2017;18:257–75.

[15] Santoro A, Balbi V, Balducci E, Pirazzini C, Rosini F, Tavano F, et al. Evidence for Sub-Haplogroup H5 of Mitochondrial DNA as a Risk Factor for Late Onset Alzheimer’s Disease. PLOS ONE 2010;5:e12037.

[16] Wu H-M, Li T, Wang Z-F, Huang S-S, Shao Z-Q, Wang K, et al. Mitochondrial DNA variants modulate genetic susceptibility to Parkinson’s disease in Han Chinese. Neurobiol Dis 2018;114:17–23.

[17] Tanaka N, Goto Y, Akanuma J, Kato M, Kinoshita T, Yamashita F, et al. Mitochondrial DNA variants in a Japanese population of patients with Alzheimer’s disease. Mitochondrion 2010;10:32–7.

[18] Tanaka M, Cabrera VM, González AM, Larruga JM, Takeyasu T, Fuku N, et al. Mitochondrial Genome Variation in Eastern Asia and the Peopling of Japan. Genome Res 2004;14:1832–50.

[19] Nagle N, Ballantyne KN, van Oven M, Tyler-Smith C, Xue Y, Wilcox S, et al. Mitochondrial DNA diversity of present-day Aboriginal Australians and implications for human evolution in Oceania. J Hum Genet 2017;62:343–53.

[20] Yonova-Doing E, Calabrese C, Gomez-Duran A, Schon K, Wei W, Karthikeyan S, et al. An atlas of mitochondrial DNA genotype–phenotype associations in the UK Biobank. Nat Genet 2021;53:982–93.

[21] Soini HK, Moilanen JS, Finnila S, Majamaa K. Mitochondrial DNA sequence variation in Finnish patients with matrilineal diabetes mellitus. BMC Res Notes 2012;5:350.

[22] Wallace DC. Mitochondria and cancer. Nat Rev Cancer 2012;12:685–98.

[23] Wei W, Schon KR, Elgar G, Orioli A, Tanguy M, Giess A, et al. Nuclear-embedded mitochondrial DNA sequences in 66,083 human genomes. Nature 2022;611:105–14.

[24] Zhou W, Karan KR, Gu W, Klein H-U, Sturm G, De Jager PL, et al. Somatic nuclear mitochondrial DNA insertions are prevalent in the human brain and accumulate over time in fibroblasts. PLoS Biol 2024;22:e3002723.

[25] Goldin E, Stahl S, Cooney AM, Kaneski CR, Gupta S, Brady RO, et al. Transfer of a mitochondrial DNA fragment to MCOLN1 causes an inherited case of mucolipidosis IV. Hum Mutat 2004;24:460–5.

[26] Turner C, Killoran C, Thomas NST, Rosenberg M, Chuzhanova NA, Johnston J, et al. Human genetic disease caused by de novo mitochondrial-nuclear DNA transfer. Hum Genet 2003;112:303–9.

[27] Ahmed ZM, Smith TN, Riazuddin S, Makishima T, Ghosh M, Bokhari S, et al. Nonsyndromic recessive deafness DFNB18 and Usher syndrome type IC are allelic mutations of USHIC. Hum Genet 2002;110:527–31.

[28] Borensztajn K, Chafa O, Alhenc-Gelas M, Salha S, Reghis A, Fischer A-M, et al. Characterization of two novel splice site mutations in human factor VII gene causing severe plasma factor VII deficiency and bleeding diathesis. Br J Haematol 2002;117:168–71.

[29] Zhuang X, Ye R, Zhou Y, Cheng MY, Cui H, Wang L, et al. Leveraging new methods for comprehensive characterization of mitochondrial DNA in esophageal squamous cell carcinoma. Genome Med 2024;16:50.

[30] Desagher S, Martinou JC. Mitochondria as the central control point of apoptosis. Trends Cell Biol 2000;10:369–77.

[31] Suen D-F, Norris KL, Youle RJ. Mitochondrial dynamics and apoptosis. Genes Dev 2008;22:1577–90.

[32] Bock FJ, Tait SWG. Mitochondria as multifaceted regulators of cell death. Nat Rev Mol Cell Biol 2020;21:85–100.

[33] Drago I, De Stefani D, Rizzuto R, Pozzan T. Mitochondrial Ca2+ uptake contributes to buffering cytoplasmic Ca2+ peaks in cardiomyocytes. Proc Natl Acad Sci 2012;109:12986–91.

[34] Cloonan SM, Choi AM. Mitochondria: sensors and mediators of innate immune receptor signaling. Curr Opin Microbiol 2013;16:327–38.

[35] Arakaki N, Nishihama T, Owaki H, Kuramoto Y, Suenaga M, Miyoshi E, et al. Dynamics of mitochondria during the cell cycle. Biol Pharm Bull 2006;29:1962–5.

[36] Finkel T, Hwang PM. The Krebs cycle meets the cell cycle: mitochondria and the G1-S transition. Proc Natl Acad Sci U S A 2009;106:11825–6.

[37] Laricchia KM, Lake NJ, Watts NA, Shand M, Haessly A, Gauthier L, et al. Mitochondrial DNA variation across 56,434 individuals in gnomAD. Genome Res 2022;32:569–82.

[38] Bolze A, Mendez F, White S, Tanudjaja F, Isaksson M, Jiang R, et al. A catalog of homoplasmic and heteroplasmic mitochondrial DNA variants in humans. bioRxiv. 2020. https://www.biorxiv.org/content/10.1101/798264

[39] Yamamoto K, Sakaue S, Matsuda K, Murakami Y, Kamatani Y, Ozono K, et al. Genetic and phenotypic landscape of the mitochondrial genome in the Japanese population. Commun Biol 2020;3:1–11.

[40] Gupta R, Kanai M, Durham TJ, Tsuo K, McCoy JG, Kotrys AV, et al. Nuclear genetic control of mtDNA copy number and heteroplasmy in humans. Nature 2023;620:839–48.

[41] Stewart JB, Chinnery PF. Extreme heterogeneity of human mitochondrial DNA from organelles to populations. Nat Rev Genet 2021;22:106–18.

[42] Zhang P, Luo H, Li Y, Wang Y, Wang J, Zheng Y, et al. NyuWa Genome resource: A deep whole-genome sequencing-based variation profile and reference panel for the Chinese population. Cell Rep 2021;37:110017.

[43] Niu Y, Teng X, Zhou H, Shi Y, Li Y, Tang Y, et al. Characterizing mobile element insertions in 5675 genomes. Nucleic Acids Res 2022;50:2493–508.

[44] Shi Y, Niu Y, Zhang P, Luo H, Liu S, Zhang S, et al. Characterization of genome-wide STR variation in 6487 human genomes. Nat Commun 2023;14:2092.

[45] Weissensteiner H, Forer L, Fendt L, Kheirkhah A, Salas A, Kronenberg F, et al. Contamination detection in sequencing studies using the mitochondrial phylogeny. Genome Res 2021;31:309–16.

[46] Ju YS, Alexandrov LB, Gerstung M, Martincorena I, Nik-Zainal S, Ramakrishna M, et al. Origins and functional consequences of somatic mitochondrial DNA mutations in human cancer. eLife 2014;3:e02935.

[47] Yuan Y, Ju YS, Kim Y, Li J, Wang Y, Yoon CJ, et al. Comprehensive molecular characterization of mitochondrial genomes in human cancers. Nat Genet 2020;52:342–52.

[48] Zhu Y, Gu X, Xu C. Mitochondrial DNA 7908–8816 region mutations in maternally inherited essential hypertensive subjects in China. BMC Med Genom 2018;11:89.

[49] Simone D, Calabrese FM, Lang M, Gasparre G, Attimonelli M. The reference human nuclear mitochondrial sequences compilation validated and implemented on the UCSC genome browser. BMC Genomics 2011;12:517.

[50] Calabrese FM, Simone D, Attimonelli M. Primates and mouse NumtS in the UCSC Genome Browser. BMC Bioinformatics 2012;13:S15.

[51] Li M, Schroeder R, Ko A, Stoneking M. Fidelity of capture-enrichment for mtDNA genome sequencing: influence of NUMTs. Nucleic Acids Research 2012;40:e137.

[52] Dayama G, Emery SB, Kidd JM, Mills RE. The genomic landscape of polymorphic human nuclear mitochondrial insertions. Nucleic Acids Research 2014;42:12640–9.

[53] McLaren W, Gil L, Hunt SE, Riat HS, Ritchie GRS, Thormann A, et al. The Ensembl Variant Effect Predictor. Genome Biol 2016;17:122.

[54] Castellana S, Biagini T, Petrizzelli F, Parca L, Panzironi N, Caputo V, et al. MitImpact 3: modeling the residue interaction network of the Respiratory Chain subunits. Nucleic Acids Res 2021;49:D1282–8.

[55] Newman NJ, Biousse V, Yu-Wai-Man P, Carelli V, Vignal-Clermont C, Montestruc F, et al. Meta-analysis of treatment outcomes for patients with m.11778G>A MT-ND4 Leber hereditary optic neuropathy. Survey of Ophthalmology 2025;70:283–95.

[56] Xie S, Zhang, Juanjuan, Sun, Jiji, Zhang, Minglian, Zhao, Fuxin, Wei, Qi-Ping, et al. Mitochondrial haplogroup D4j specific variant m.11696G > a(MT-ND4) may increase the penetrance and expressivity of the LHON-associated m.11778G > a mutation in Chinese pedigrees. Mitochondrial DNA Part A 2017;28:434–41.

[57] Sonney S, Leipzig J, Lott MT, Zhang S, Procaccio V, Wallace DC, et al. Predicting the pathogenicity of novel variants in mitochondrial tRNA with MitoTIP. PLoS Comput Biol 2017;13:e1005867.

[58] McManus MJ, Picard M, Chen H-W, De Haas HJ, Potluri P, Leipzig J, et al. Mitochondrial DNA Variation Dictates Expressivity and Progression of Nuclear DNA Mutations Causing Cardiomyopathy. Cell Metab 2019;29:78–90.e5.

[59] Pickett SJ, Deen D, Pyle A, Santibanez-Koref M, Hudson G. Interactions between nuclear and mitochondrial SNPs and Parkinson’s disease risk. Mitochondrion 2022;63:85–8.

[60] Andrews SJ, Fulton-Howard B, Patterson C, McFall GP, Gross A, Michaelis EK, et al. Mitonuclear interactions influence Alzheimer’s disease risk. Neurobiol Aging 2020;87:138.e7–138.e14.

[61] Xia C, Pickett SJ, Liewald DCM, Weiss A, Hudson G, Hill WD. The contributions of mitochondrial and nuclear mitochondrial genetic variation to neuroticism. Nat Commun 2023;14:3146.

[62] van Oven M, Kayser M. Updated comprehensive phylogenetic tree of global human mitochondrial DNA variation. Hum Mutat 2009;30:E386–94.

[63] Jiang P, Zhu T, Liu J, Tao X, Xue Z, Tao Y, et al. Mitochondrial DNA variant spectrum and the association with chronic tic disorders. European Journal of Neurology 2022;29:3187–96.

[64] Ludwig-Słomczyńska AH, Seweryn MT, Kapusta P, Pitera E, Handelman SK, Mantaj U, et al. Mitochondrial GWAS and association of nuclear – mitochondrial epistasis with BMI in T1DM patients. BMC Medical Genomics 2020;13:97.

[65] Modi A, Lancioni H, Cardinali I, Capodiferro MR, Rambaldi Migliore N, Hussein A, et al. The mitogenome portrait of Umbria in Central Italy as depicted by contemporary inhabitants and pre-Roman remains. Sci Rep 2020;10:10700.

[66] Guichard JL, Kane MS, Grenett M, Sandel M, Benavides GA, Bradley WE, et al. Mitochondrial haplotype modulates genome expression and mitochondrial structure/function in cardiomyocytes following volume overload. American Journal of Physiology-Heart and Circulatory Physiology 2023.

[67] Weissensteiner H, Pacher D, Kloss-Brandstätter A, Forer L, Specht G, Bandelt H-J, et al. HaploGrep 2: mitochondrial haplogroup classification in the era of high-throughput sequencing. Nucleic Acids Res 2016;44:W58–63.

[68] Behar DM, van Oven M, Rosset S, Metspalu M, Loogväli E-L, Silva NM, et al. A “Copernican” Reassessment of the Human Mitochondrial DNA Tree from its Root. Am J Hum Genet 2012;90:675–84.

[69] Xue L, Moreira JD, Smith KK, Fetterman JL. The Mighty NUMT: Mitochondrial DNA Flexing Its Code in the Nuclear Genome. Biomolecules 2023;13:753.

[70] Blok MJ, Spruijt L, Coo IFM de, Schoonderwoerd K, Hendrickx A, Smeets HJ. Mutations in the ND5 subunit of complex I of the mitochondrial DNA are a frequent cause of oxidative phosphorylation disease. J Med Genet 2007;44:e74–e74.

[71] Tang S, Huang T. Characterization of mitochondrial DNA heteroplasmy using a parallel sequencing system. Biotechniques 2010;48:287–96.

[72] Huang T. Next generation sequencing to characterize mitochondrial genomic DNA heteroplasmy. Curr Protoc Hum Genet 2011;Chapter 19:19.8.1–19.8.12.

[73] Xu W, Ghosh S, Comhair SAA, Asosingh K, Janocha AJ, Mavrakis DA, et al. Increased mitochondrial arginine metabolism supports bioenergetics in asthma. J Clin Investig 2016;126:2465–81.

[74] Kang E, Wu J, Gutierrez NM, Koski A, Tippner-Hedges R, Agaronyan K, et al. Mitochondrial replacement in human oocytes carrying pathogenic mitochondrial DNA mutations. Nature 2016;540:270–5.

[75] Gorman GS, Schaefer AM, Ng Y, Gomez N, Blakely EL, Alston CL, et al. Prevalence of nuclear and mitochondrial DNA mutations related to adult mitochondrial disease. Ann Neurol 2015;77:753–9.

[76] Hazkani-Covo E, Zeller RM, Martin W. Molecular Poltergeists: Mitochondrial DNA Copies (numts) in Sequenced Nuclear Genomes. PLOS Genet 2010;6:e1000834.

[77] Chen J-M, Chuzhanova N, Stenson PD, Férec C, Cooper DN. Meta-analysis of gross insertions causing human genetic disease: novel mutational mechanisms and the role of replication slippage. Hum Mutat 2005;25:207–21.

[78] Willsey AJ, Fernandez TV, Yu D, King RA, Dietrich A, Xing J, et al. De Novo Coding Variants Are Strongly Associated with Tourette Disorder. Neuron 2017;94:486–499.e9.

[79] Kraja AT, Liu C, Fetterman JL, Graff M, Have CT, Gu C, et al. Associations of Mitochondrial and Nuclear Mitochondrial Variants and Genes with Seven Metabolic Traits. The American Journal of Human Genetics 2019;104:112–38.

[80] Sukhorukov VN, Orekhov AN. Molecular Aspects of Inflammation and Lipid Metabolism in Health and Disease: The Role of the Mitochondria. International Journal of Molecular Sciences 2024;25:6299.

[81] Liu X, Sun X, Zhang Y, Jiang W, Lai M, Wiggins KL, et al. Association Between Whole Blood–Derived Mitochondrial DNA Copy Number, Low-Density Lipoprotein Cholesterol, and Cardiovascular Disease Risk. Journal of the American Heart Association 2023.

[82] Fedotova EI, Berezhnov AV, Popov DY, Shitikova EY, Vinokurov AY. The Role of mtDNA Mutations in Atherosclerosis: The Influence of Mitochondrial Dysfunction on Macrophage Polarization. International Journal of Molecular Sciences 2025;26:1019.

[83] Boone C, Lewis SC. Bridging lipid metabolism and mitochondrial genome maintenance. Journal of Biological Chemistry 2024;300.

[84] He Y, Wu J, Dressman DC, Iacobuzio-Donahue C, Markowitz SD, Velculescu VE, et al. Heteroplasmic mitochondrial DNA mutations in normal and tumour cells. Nature 2010;464:610–4.

[85] Li H, Durbin R. Fast and accurate long-read alignment with Burrows-Wheeler transform. Bioinformatics 2010;26:589–95.

[86] DePristo MA, Banks E, Poplin R, Garimella KV, Maguire JR, Hartl C, et al. A framework for variation discovery and genotyping using next-generation DNA sequencing data. Nat Genet 2011;43:491–8.

[87] Poplin R, Ruano-Rubio V, DePristo MA, Fennell TJ, Carneiro MO, Van Der Auwera GA, et al. Scaling accurate genetic variant discovery to tens of thousands of samples 2017.

[88] Zhang F, Flickinger M, Taliun SAG, InPSYght Psychiatric Genetics Consortium, Abecasis GR, Scott LJ, et al. Ancestry-agnostic estimation of DNA sample contamination from sequence reads. Genome Res 2020;30:185–94.

[89] Krzywinski M, Schein J, Birol I, Connors J, Gascoyne R, Horsman D, et al. Circos: an information aesthetic for comparative genomics. Genome Res 2009;19:1639–45.

[90] Lott MT, Leipzig JN, Derbeneva O, Xie HM, Chalkia D, Sarmady M, et al. mtDNA Variation and Analysis Using Mitomap and Mitomaster. Curr Protoc Bioinformatics 2013;44:1.23.1-26.

[91] Quinlan AR, Hall IM. BEDTools: a flexible suite of utilities for comparing genomic features. Bioinformatics 2010;26:841–2.

[92] Navarro Gonzalez J, Zweig AS, Speir ML, Schmelter D, Rosenbloom KR, Raney BJ, et al. The UCSC Genome Browser database: 2021 update. Nucleic Acids Res 2021;49:D1046–57.

