## Supplemental Materials for "Mitochondrial Genome Variants and Nuclear Mitochondrial DNA Segments in 7331 Individuals from NyuWa and 1KGP"

### Supplemental Figures

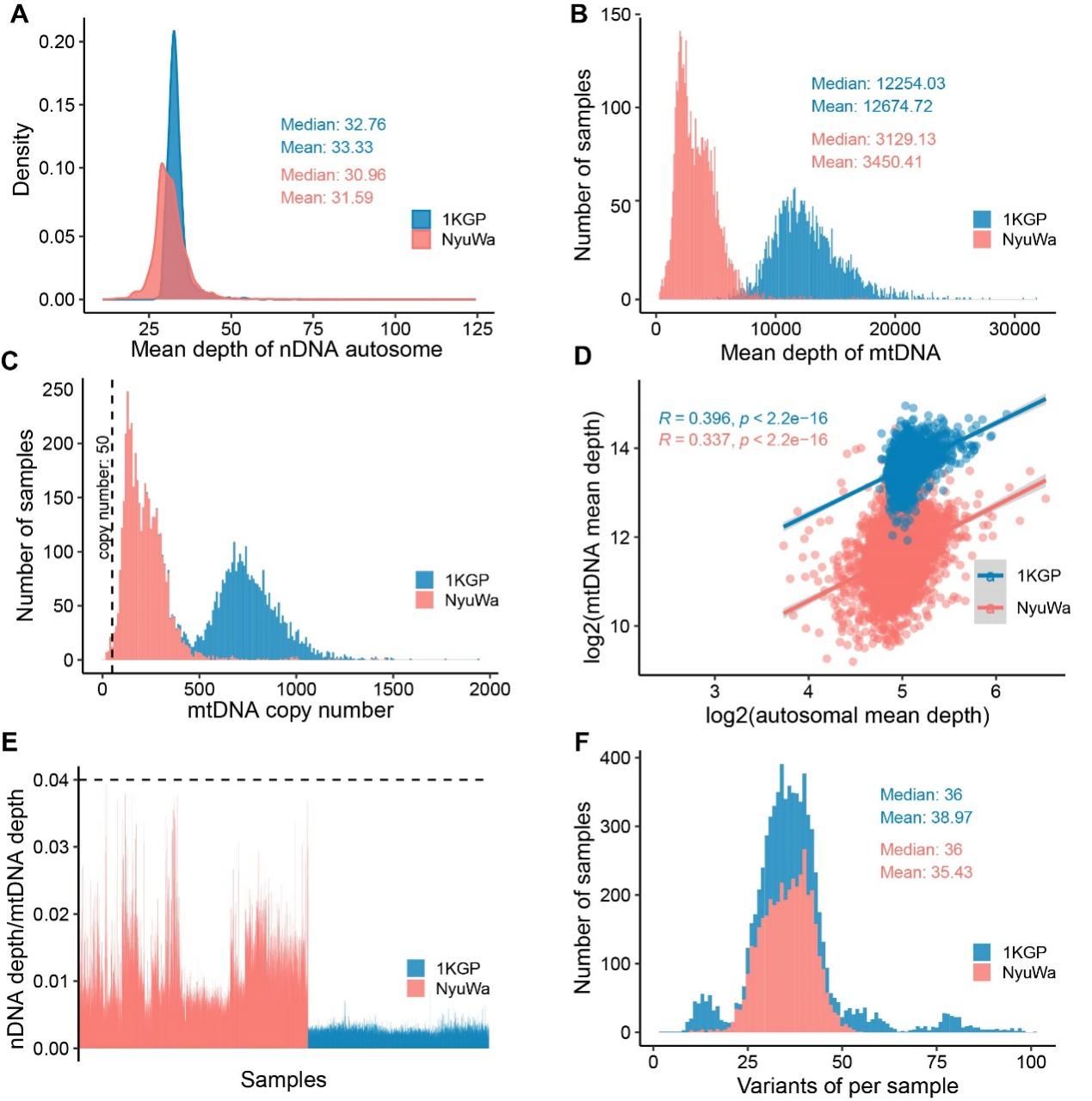

**Figure S1 High-quality mtDNA variants**

**A**. Distribution of nDNA mean depth in individuals (n = 7331). **B.** Distribution of mtDNA mean depth in individuals(n = 7331). **C.** Distribution of copy number of mtDNA in this cohort. **D.** Correlation of autosomal mean depth and mtDNA mean depth. **E.** The ratio of nDNA to mtDNA depth in all samples of this cohort. **F.** Distribution of high-quality mtDNA variants for each sample in this study.

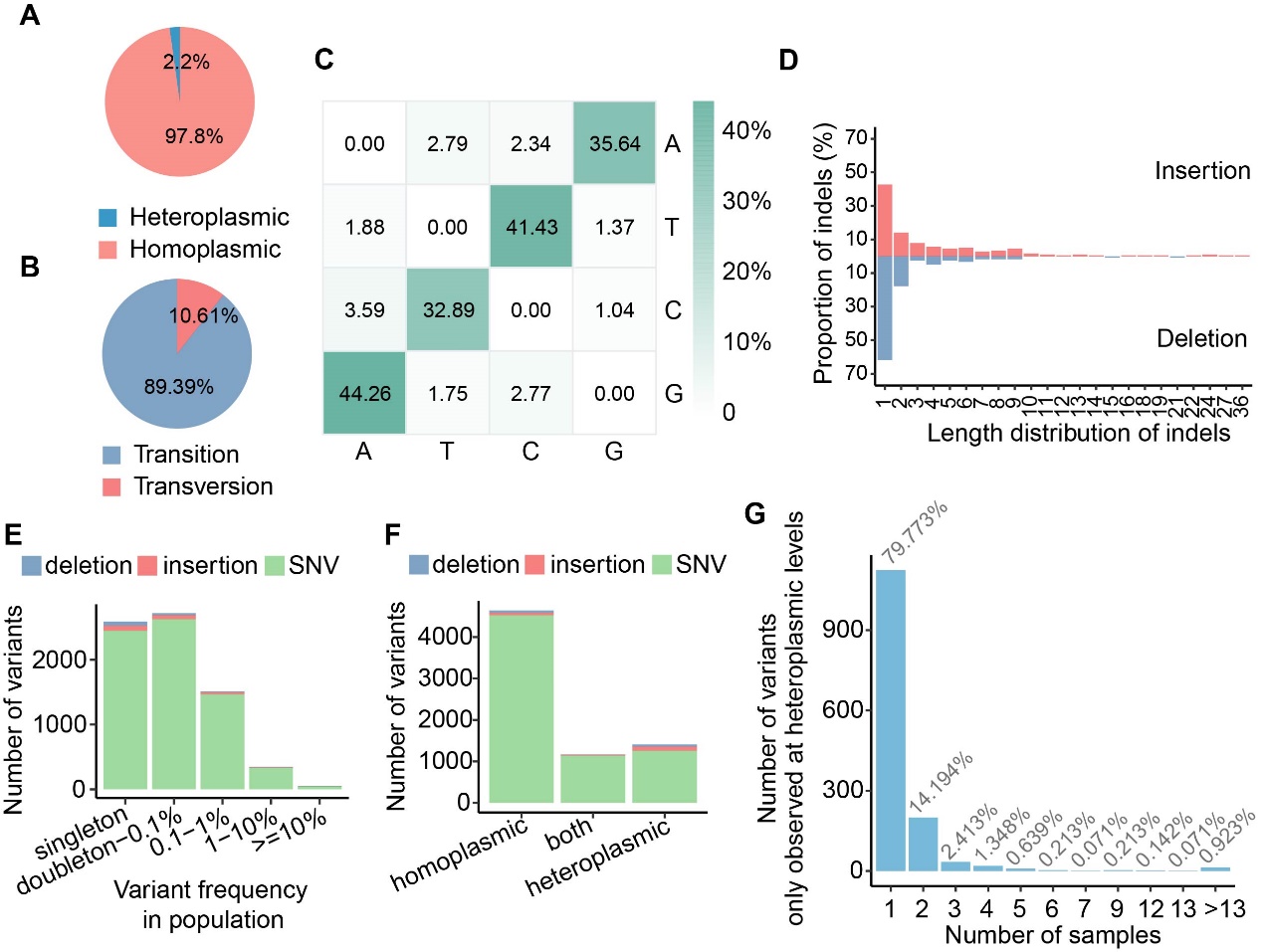

**Figure S2 Characteristics of mtDNA variants**

**A.** Proportion of homoplasmic and heteroplasmic variants in total mtDNA variants. **B.** Proportion of the transition and transversion SNVs in this study. **C.** Proportion heatmap of transition and transversion SNVs. The horizontal axis represents alt bases, and the vertical axis represents ref bases. The denominators of each row of the matrix are the total number of A, T, C, and G on the mitochondrial reference genome. **D.** Length distribution of mtDNA indels in this study. **E.** Variant types of non-redundant high-quality mtDNA variants detected at different variant frequencies in this study. **F.** Variant types of non-redundant high-quality mtDNA variants detected at different heteroplamic levels (homoplasmic, heteroplasmic, and both). **G**. Distribution of variants only observed at heteroplasmic levels in this study.

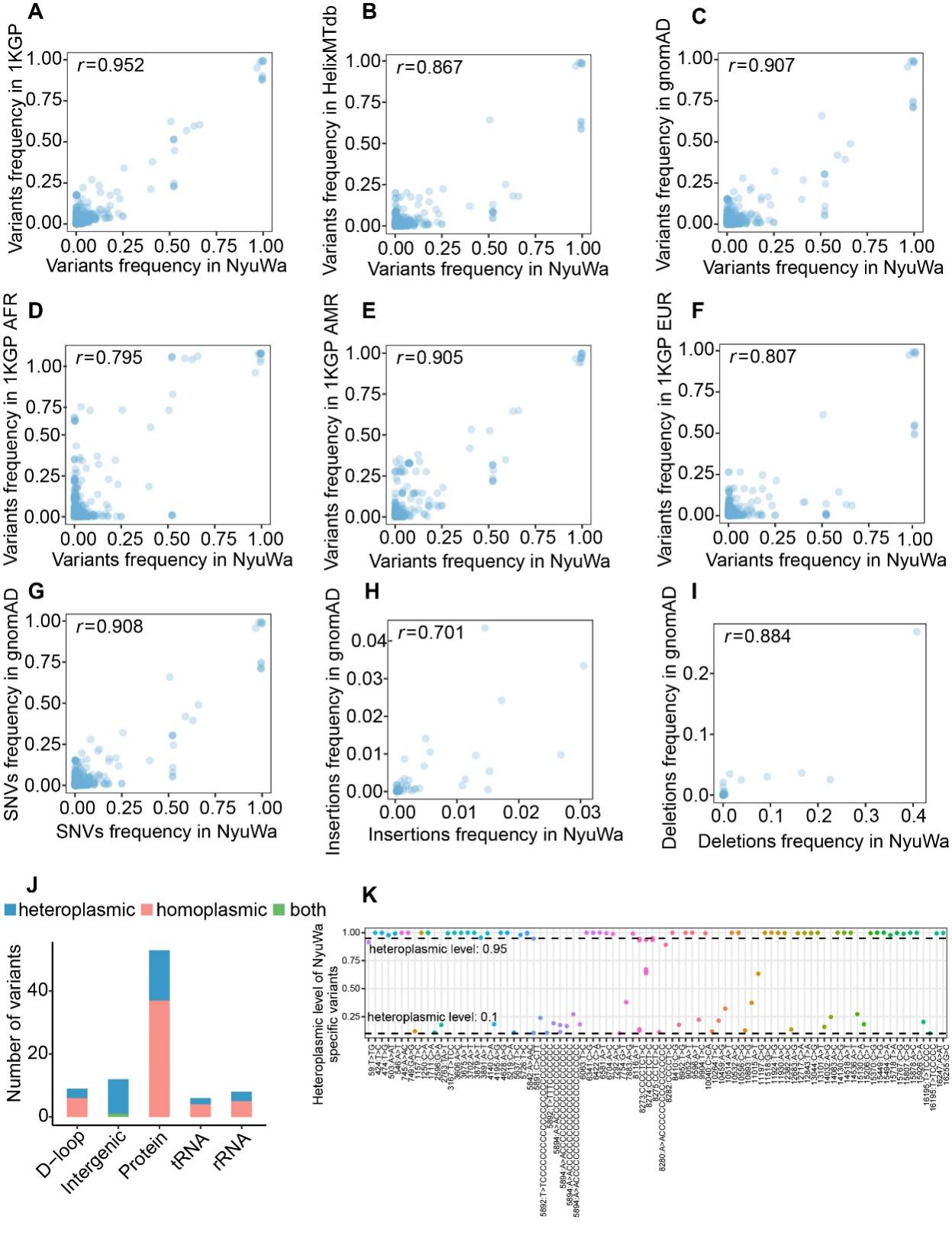

**Figure S3 Comparison of mtDNA variants in NyuWa with other resources**

**A**. Frequency of variants between NyuWa and 1KGP. Pearson correlation coefficients were shown in the figure. **B**. Frequency of variants between NyuWa and HelixMTdb. **C**. Frequency of variants between NyuWa and gnomAD. **D.** Frequency of variants between NyuWa and AFR of 1KGP. AFR, African. **E.** Frequency of variants between NyuWa and AMR of 1KGP. AMR, American. **F.** Frequency of variants between NyuWa and EUR of 1KGP. EUR, European. **G.** Frequency of SNVs between NyuWa and gnomAD. **H.** Frequency of insertions between NyuWa and gnomAD. **I.** Frequency of deletions between NyuWa and gnomAD. **J.** Number of NyuWa-specific variants at different heteroplasmic levels in mtDNA regions. **K.** Shown all heteroplasmic levels of 88 NyuWa-specific variants in this cohort. Different colors represent different variants.

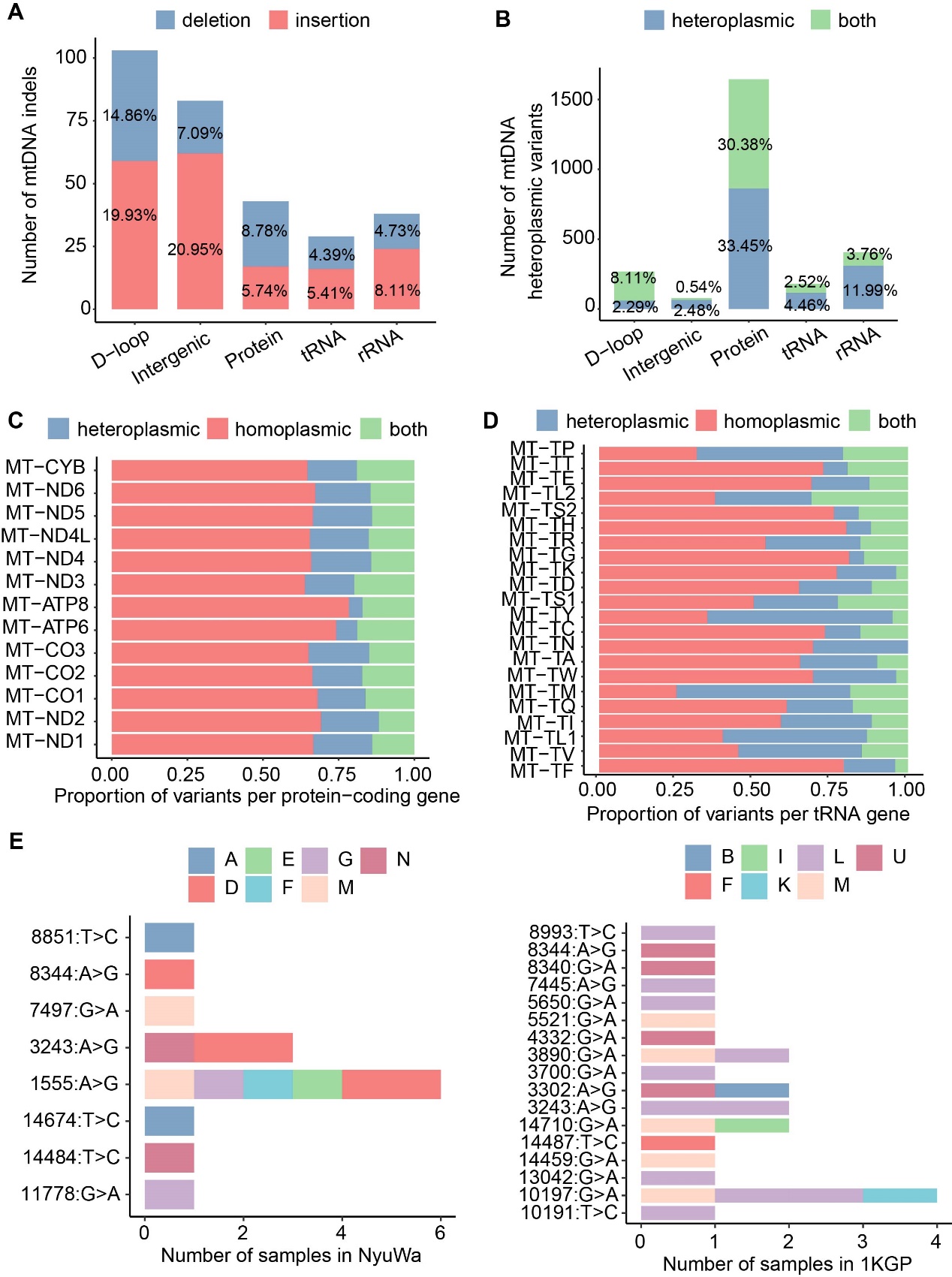

**Figure S4 Predicted functions and pathogenicity of mtDNA variants**

**A.** Number of indels in mtDNA regions. **B.** Number of mtDNA heteroplasmic variants in mtDNA regions. **C.** Proportion of variants (homoplasmic, heteroplasmic, and both) per protein-coding gene. **D.** Proportion of variants (homoplasmic, heteroplasmic, and both) per tRNA gene. **E.** Known pathogenic variants observed in this study along with the haplogroups.

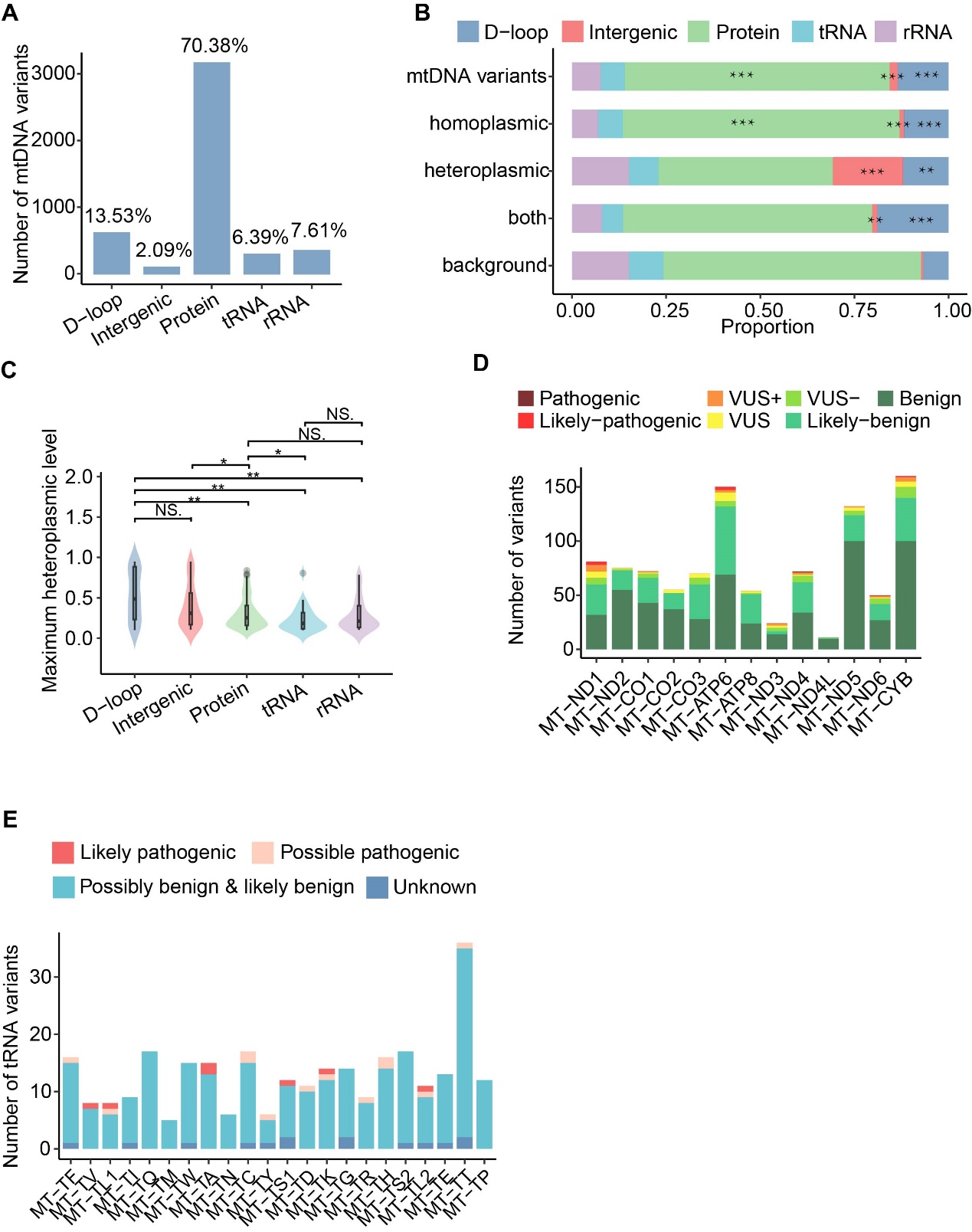

**Figure S5 Predicted functions and pathogenicity of mtDNA variants in NyuWa**

**A.** Number of mtDNA variants annotated to the mtDNA regions. **B.** Proportion of variant annotations in mtDNA regions. mtDNA variants for total variants. Homoplasmic for variants observed only at homoplasmic levels. Heteroplasmic for variants observed only at heteroplasmic levels. Both for variants observed both at homoplasmic and heteroplasmic levels. Background for the mtDNA genome. ** for *P* <= 0.01, *** for *P* <= 0.001 based on one-tailed Hypergeometric test. **C.** The distribution of the maximum heteroplasmic levels of mtDNA variants in mtDNA regions. *** for *P* <= 0.001, ** for *P* <= 0.01, NS. for no significance based on one-tailed Wilcoxon test. **D.** Number of missense variants in protein-coding genes. APOGEE 2 refers to five pathogenicity classes: benign, likely-benign, VUS, likely-pathogenic, and pathogenic. VUS+ means closer to the likely pathogenic threshold. VUS- means closer to the likely benign threshold. **E.** Number of variants in tRNA genes. The severity of variants was defined by MitoTIP. Likely pathogenic: Variants with high pathogenicity scores, indicating a strong likelihood of causing disease. Possible pathogenic: Variants with moderate pathogenicity scores that may be associated with disease, suggesting a potential but less certain disease-causing effect. Possibly benign & likely benign: Variants with low pathogenicity scores, unlikely to cause disease. Unknown: Variants that do not have a MitoTIP percentile score.

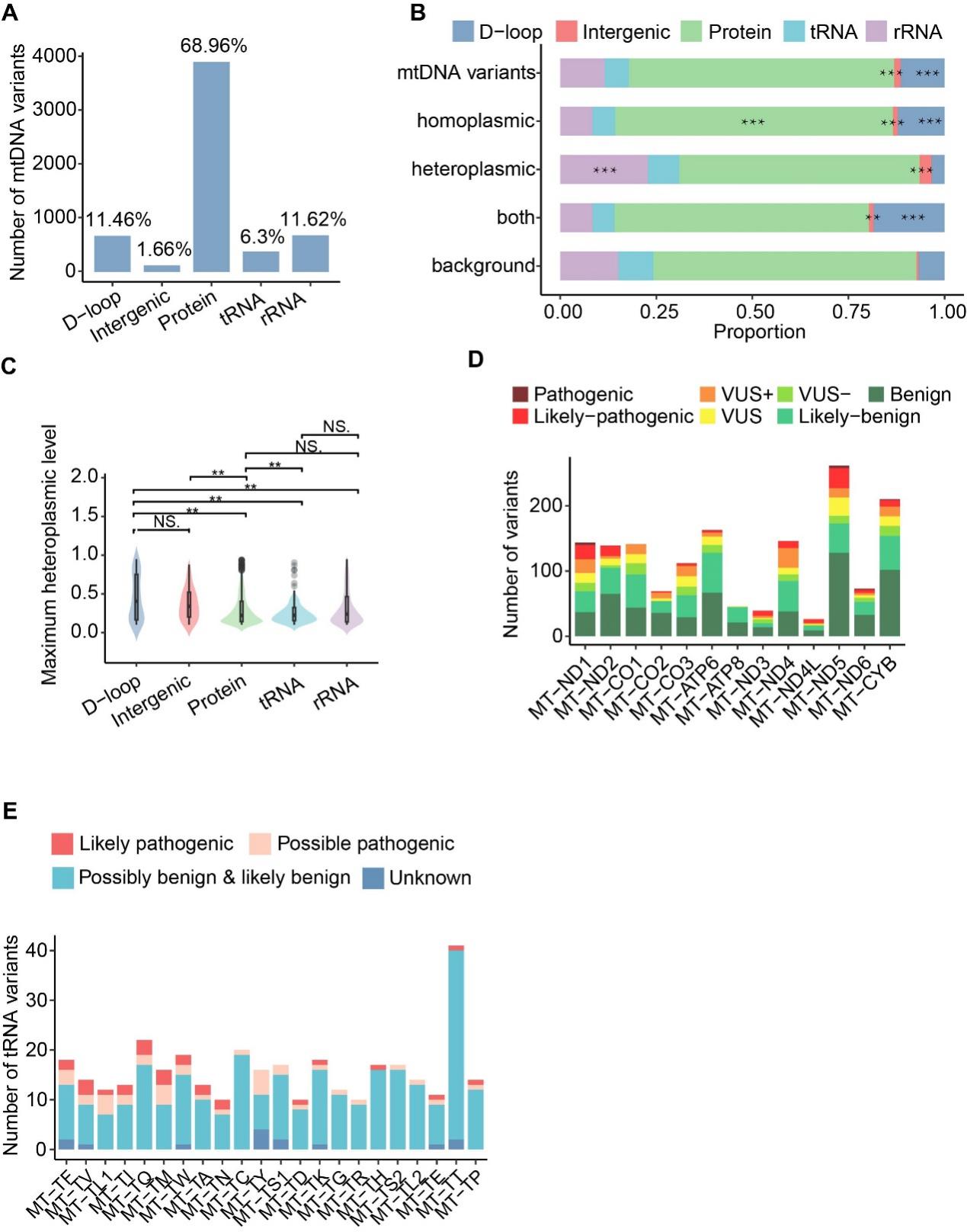

**Figure S6 Predicted functions and pathogenicity of mtDNA variants in 1KGP**

**A.** Number of mtDNA variants annotated to the mtDNA regions. **B.** Proportion of variant annotations in mtDNA regions. mtDNA variants for total variants. Homoplasmic for variants observed only at homoplasmic levels. Heteroplasmic for variants observed only at heteroplasmic levels. Both for variants observed both at homoplasmic and heteroplasmic levels. Background for the mtDNA genome. ** for *P* <= 0.01, *** for *P* <= 0.001 based on one-tailed Hypergeometric test. **C.** The distribution of the maximum heteroplasmic levels of mtDNA variants in mtDNA regions. *** for *P* <= 0.001, ** for *P* <= 0.01, NS. for no significance based on one-tailed Wilcoxon test. **D.** Number of missense variants in protein-coding genes. APOGEE 2 refers to five pathogenicity classes: benign, likely-benign, VUS, likely-pathogenic, and pathogenic. VUS+ means closer to the likely pathogenic threshold. VUS- means closer to the likely benign threshold. **E.** Number of variants in tRNA genes. The severity of variants was defined by MitoTIP. Likely pathogenic: Variants with high pathogenicity scores, indicating a strong likelihood of causing disease. Possible pathogenic: Variants with moderate pathogenicity scores that may be associated with disease, suggesting a potential but less certain disease-causing effect. Possibly benign & likely benign: Variants with low pathogenicity scores, unlikely to cause disease. Unknown: Variants that do not have a MitoTIP percentile score.

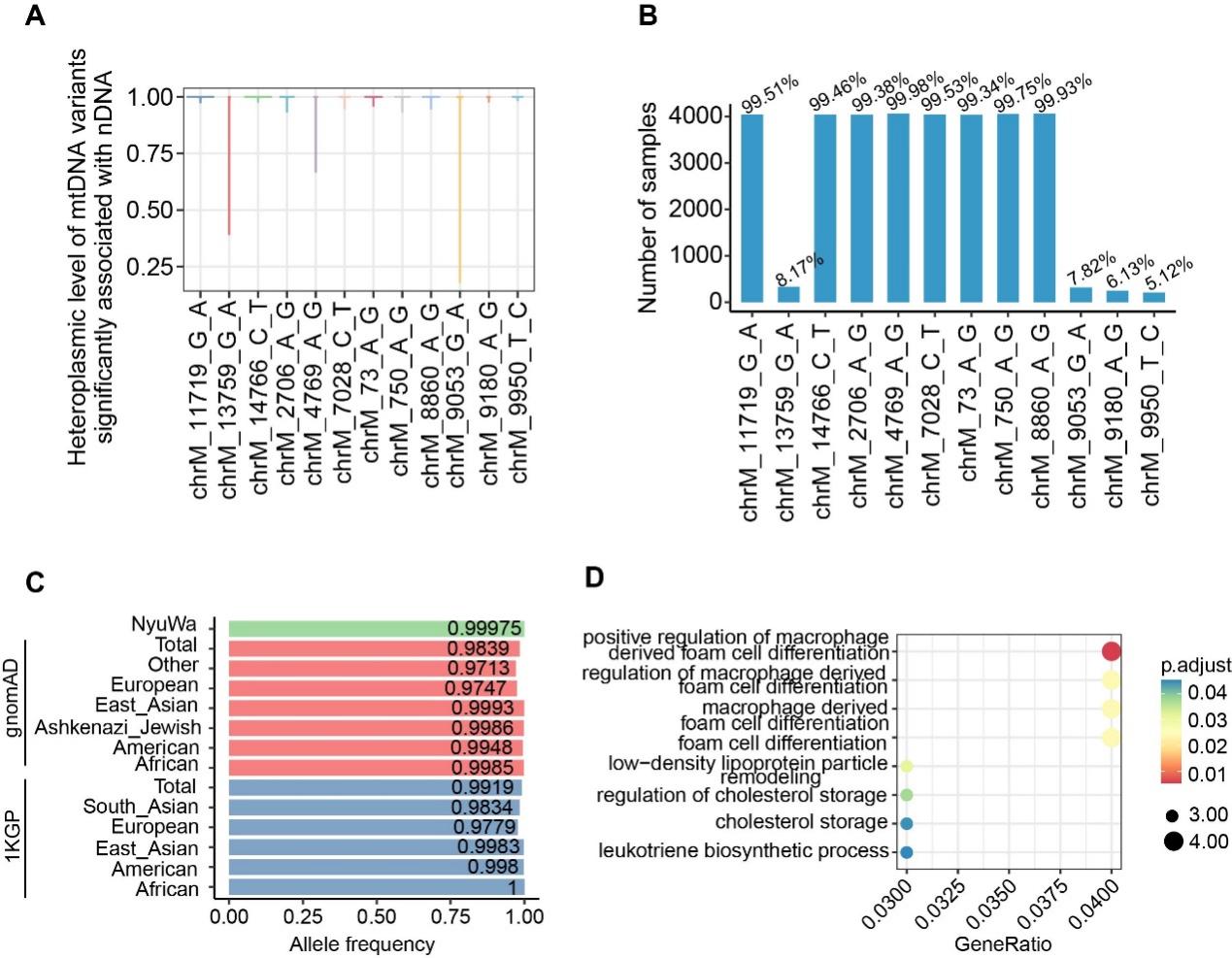

**Figure S7 mtDNA variants significantly associated with nDNA variants**

**A.** Heteroplasmic levels of 12 mtDNA variants associated with nDNA variants. **B**. Number of samples containing the 12 mtDNA variants in this study. **C**. Population/Variant frequencies of mtDNA variant, m.4769A>G, in NyuWa, gnomAD, and 1KGP. The term "Total" refers to the 1KGP or genomAD entire dataset. The category 'Other' encompasses samples that do not belong to the listed populations (European, East Asian, Ashkenazi Jewish, American, and African) and includes additional populations not specifically detailed in the figure. D. Enriched gene ontology (GO) pathways for the genes located within a 25kb upstream and downstream range of the 199 nDNA variants.

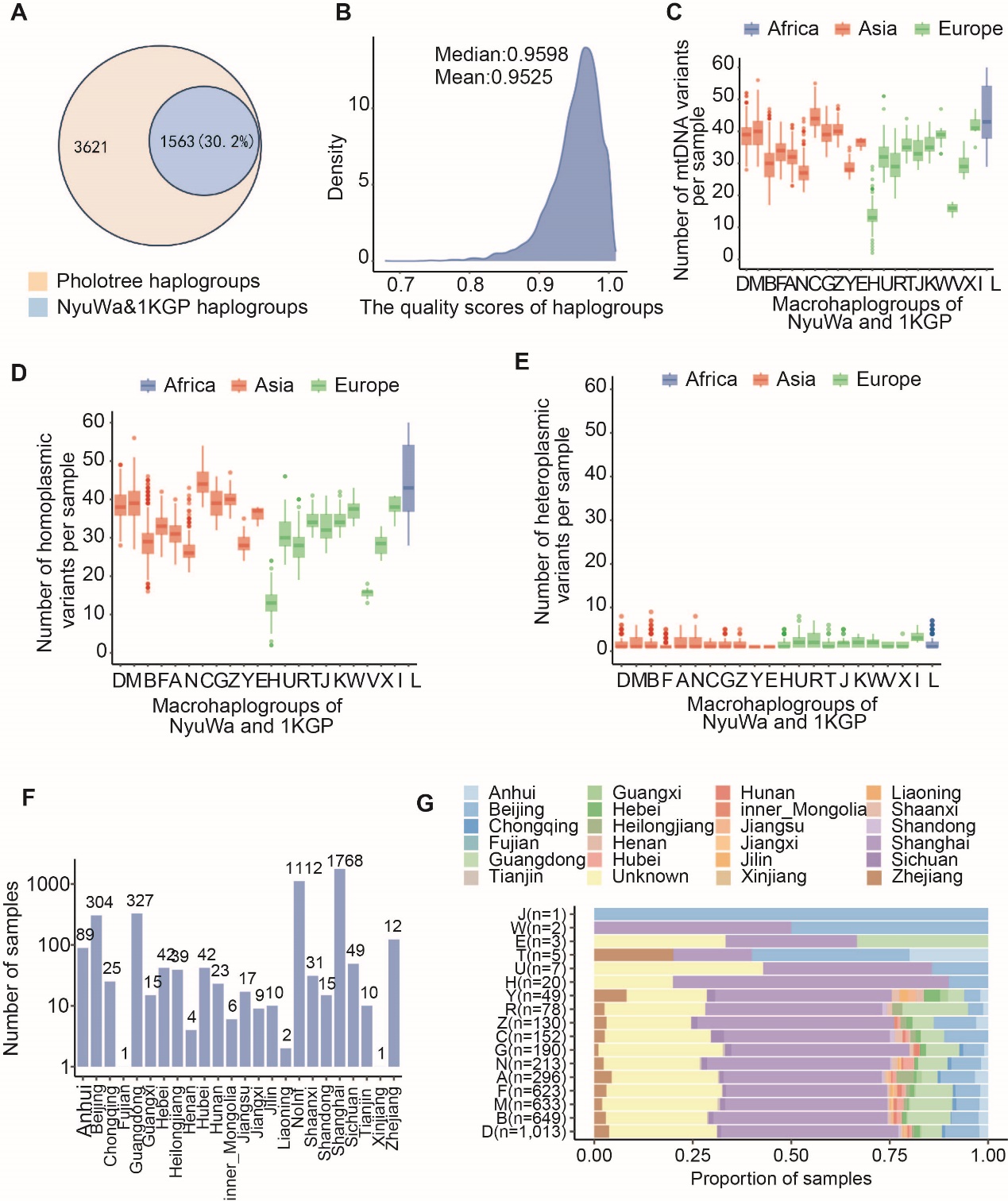

**Figure S8 Haplogroup composition in this study (related to Figure 5)**

**A.** Venn diagram of overlapping mtDNA haplogroups in this study and Phylotree. **B.** The distribution of the quality scores for haplogroups based on Haplogrep software. Median and mean quality scores were shown in figure. **C.** Number of mtDNA variants across samples within each macrohaplogroup. Colors indicated haplogroups associated with Africa, Asian and European, according to the population division of MITOMAP. **D.** The number of homoplasmic variants across samples within each macrohaplogroup. Colors indicated haplogroups associated with Africa, Asian and European, according to the population division of MITOMAP. **E.** The number of heteroplamsic variants across samples within each macrohaplogroup. **F.** The number of samples in each administrative region in NyuWa. **G.** The composition of sample origins for each macrohaplogroup.

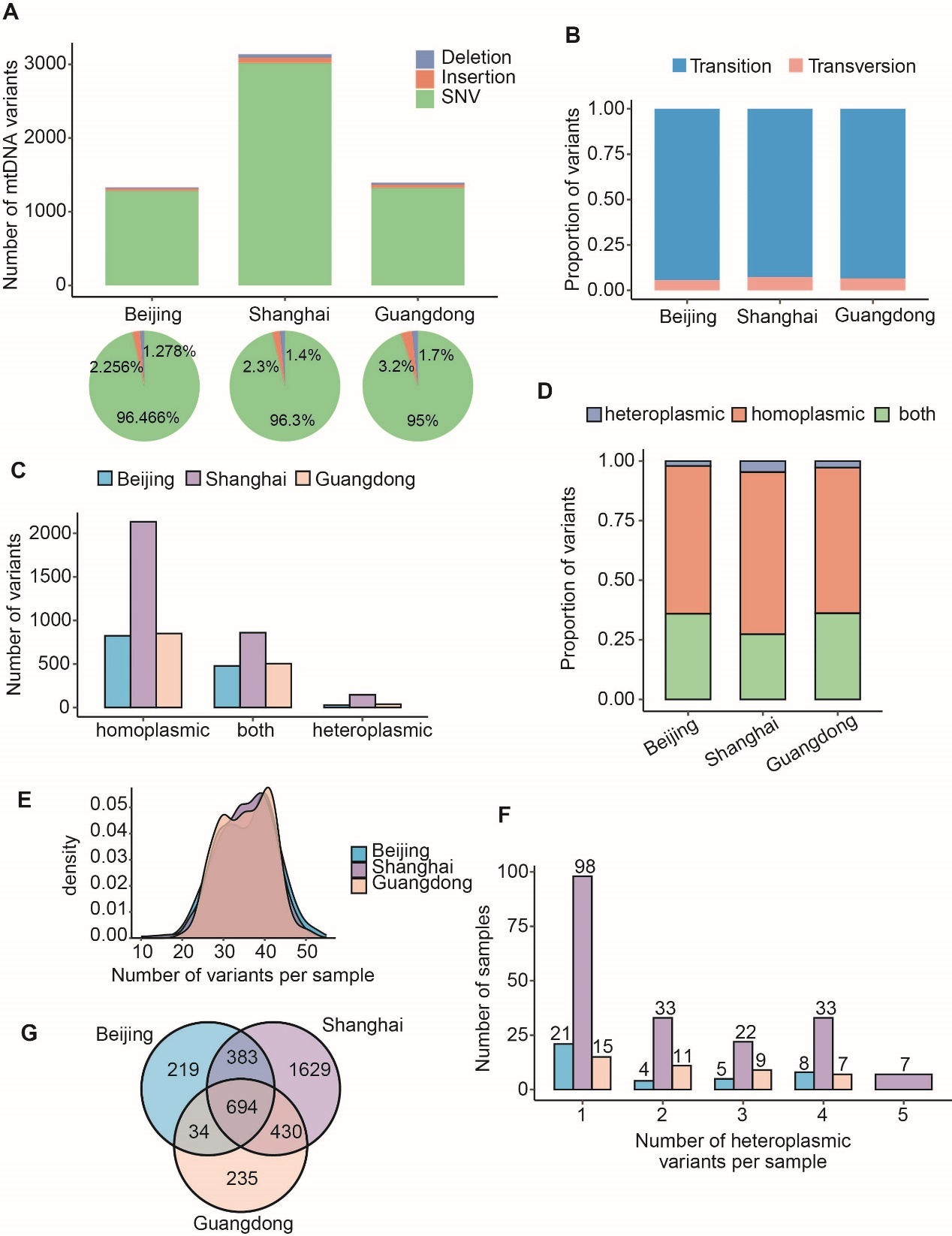

**Figure S9 Characteristics of mtDNA variants in Beijing, Shanghai and Guangdong**

**A.** Variant types (SNVs, deletions, insertions) of mtDNA variants detected at different administrative region (Beijing, Shanghai and Guangdong). **B.** Proportion of the transition and transversion SNVs in Beijing, Shanghai and Guangdong. **C.** Number of variants (homoplasmic, heteroplasmic, and both) in Beijing, Shanghai and Guangdong. Considering that Beijing and Guangdong only have about 300 samples, the heteroplasmic level classification of the variants were not re-performed here. **D.** Proportion of variants (homoplasmic, heteroplasmic, and both) in Beijing, Shanghai and Guangdong. **E.** Distribution of mtDNA variants per sample in Beijing, Shanghai and Guangdong. **F.** Number of heteroplasmic variants per sample in Beijing, Shanghai and Guangdong. **G.** Venn diagram of mtDNA variants distribution among Beijing, Shanghai, and Guangdong.

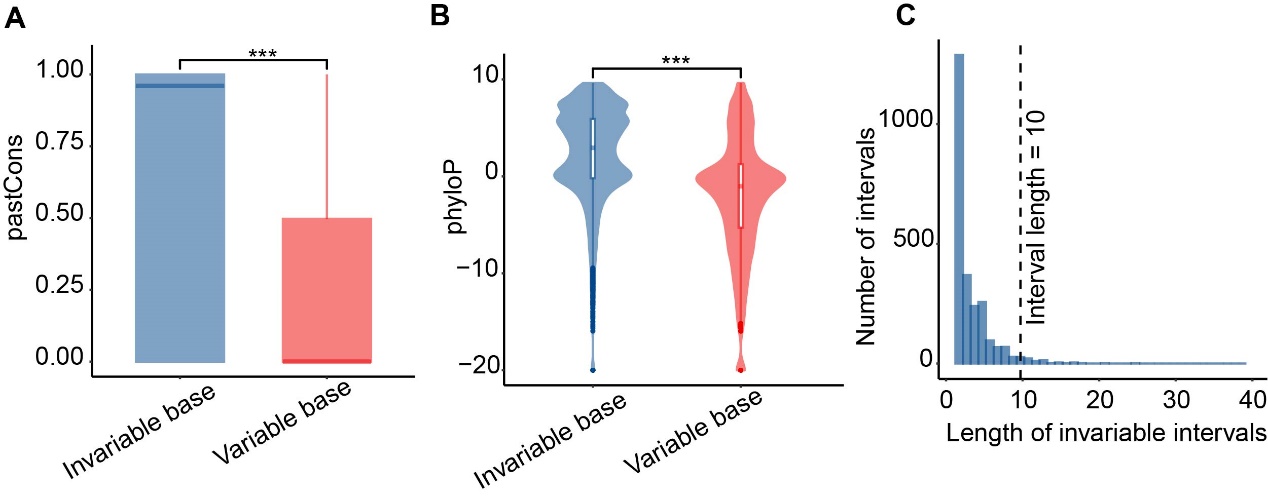

**Figure S10 Invariable segments in mtDNA**

**A.** phastCons scores of invariable and variable bases in mtDNA of this study. *** indicating *P* < 0.001 based on one-tailed Wilcoxon test. **B.** phyloP scores of invariable and variable bases in mtDNA of this study. *** indicating *P* < 0.001 based on one-tailed Wilcoxon test. **C.** The length distribution of the invariable intervals with length > 1 nt.

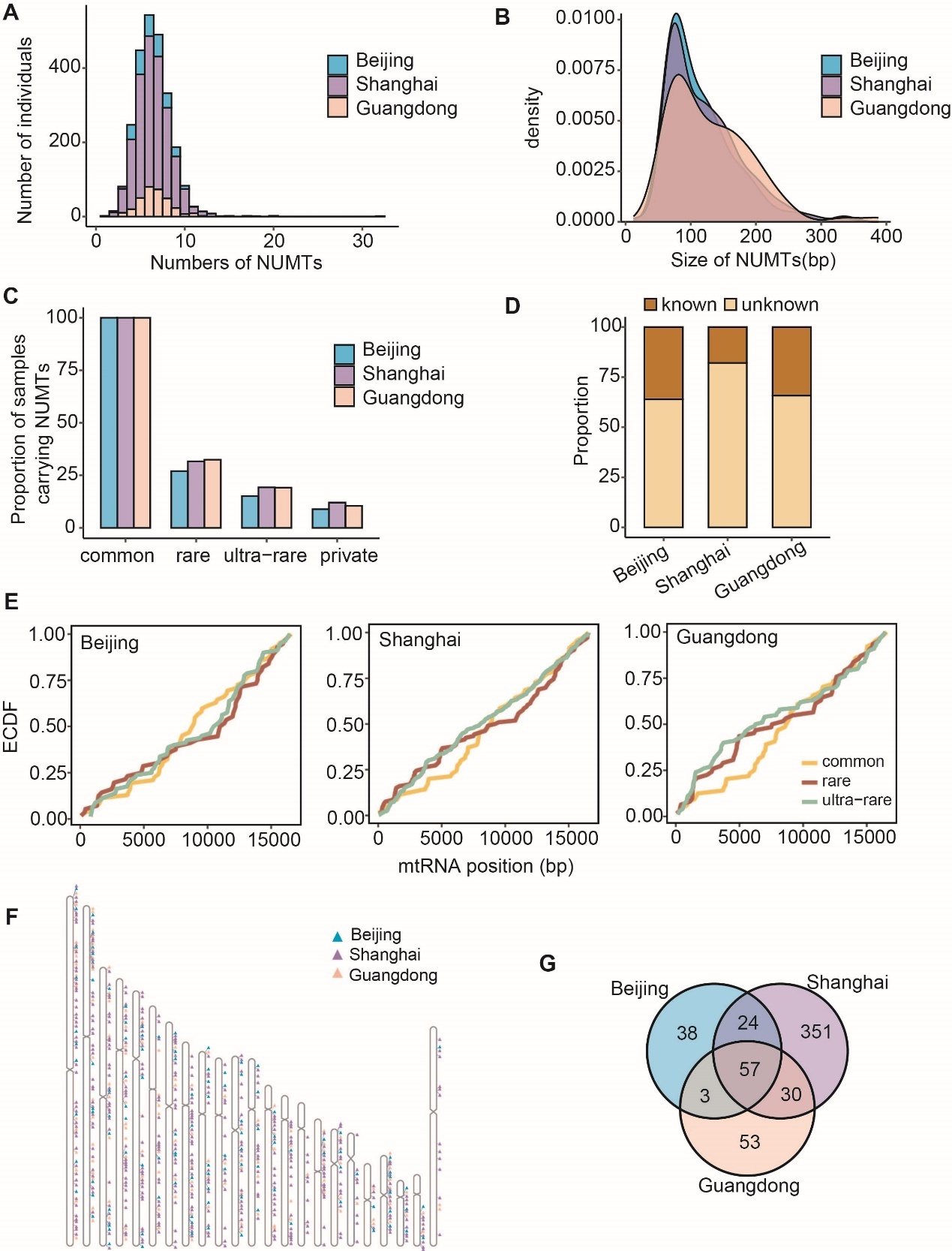

**Figure S11 Characteristics of NUMTs in Beijing, Shanghai and Guangdong**

**A.** The number distribution of NUMTs in Beijing, Shanghai and Guangdong. **B.** Size distribution of NUMTs less than 400 bp in Beijing, Shanghai and Guangdong. **C.** Proportion of samples carrying NUMTs by population frequency. Considering that Beijing and Guangdong only have about 300 samples, population frequencies of NUMTs were not recalculated here. **D.** Proportion of reported (known) and newly (unknown) identified NUMTs in Beijing, Shanghai and Guangdong. **E.** Empirical cumulative distribution function (ECDF) plot of NUMT mitochondrial breakpoints in Beijing, Shanghai and Guangdong. **F.** Chromosome map of chromosomal locations NUMTs inserted. **G.** Venn diagram of NUMTs distribution among Beijing, Shanghai, and Guangdong.
